# Biomechanical Regulation of Breast Cancer Metastasis and Progression

**DOI:** 10.1101/2020.01.21.914242

**Authors:** Adrianne Spencer, Andrew D. Sligar, Daniel Chavarria, Jason Lee, Darshil Choksi, Nikita P. Patil, HooWon Lee, Austin P. Veith, William J. Riley, Shubh Desai, Ali Abbaspour, Rohan Singeetham, Aaron B. Baker

## Abstract

Physical activity has been consistently linked to decreased incidence of breast cancer and a substantial increase in the length of survival of patients with breast cancer. However, the understanding of how applied physical forces directly regulate breast cancer remains limited. We investigated the role of mechanical forces in altering the chemoresistance, proliferation and metastasis of breast cancer cells. We found that applied mechanical tension can dramatically alter gene expression in breast cancer cells, leading to decreased proliferation, increased resistance to chemotherapeutic treatment and enhanced adhesion to inflamed endothelial cells and collagen I under fluidic shear stress. A mechanistic analysis of the pathways involved in these effects supported a complex signaling network that included Abl1, Lck, Jak2 and PI3K to regulate pro-survival signaling and enhancement of adhesion under flow. Studies using mouse xenograft models demonstrated reduced proliferation of breast cancer cells with orthotopic implantation and increased metastasis to the skull when the cancer cells were treated with mechanical load. Using high throughput mechanobiological screens we identified pathways that could be targeted to reduce the effects of load on metastasis and found that the effects of mechanical load on bone colonization could be reduced through treatment with a PI3Kγ inhibitor.

## Introduction

Cancer cells are subjected to a complex mechanical microenvironment that includes extracellular matrix compliance, alterations in local mechanical stress due to tumor mass expansion, and applied force from interactions with surrounding organs and body motion. Biophysical forces are emerging as powerful regulators of cancer growth, quiescence and metastasis but our understanding of the mechanisms of the biomechanical regulation of tumor biology remains very limited (1–3). While there is an increased appreciation of the role of tumor stiffness in regulating cancer biology (4–6), the role of externally applied forces in the different stages of cancer progression is poorly understood. In normal breast tissues, there is a distribution of tension and compressive forces that are applied dynamically to the growing tumor mass (7).

Several studies have also indicated that in rapidly growing tumors there is the development of compressive forces inside the tumor with tensional forces at the outer surface of growing tumor mass (8). In prior studies, these compressive forces were found to increase the invasive phenotype of the cells (8). Tensional forces are present in several regions of the growing cancer mass including the periphery of the tumor due to rapid tumor expansion, at the tumor/vasculature interface, and due to contraction of the skin/muscles during the motion of the body in exercise or daily activities (9, 10). Biomechanical models of the breast suggest that the tissues are subjected to cyclic forces with peaks of approximately 5 to 15 N around 5,000 times per day from walking alone (11). The modulus of glandular tissue in the breast ranges between 7.5 to 66 kPa, implying that the breast is subjected to cyclic mechanical strains over a range from 1% to 25% strain during daily activity (11). Thus, there are a rich variety of mechanical conditions within the breast tissue normally and these forces act on tumors that form within the tissue, potentially altering the course of the disease. Studies of the role of applied forces in regulating breast cancer support that compressive forces may enhance invasiveness (8) but may also suppress proliferation (12, 13). However, the understanding of how applied mechanical forces alter breast cancer progression and metastasis remain poorly understood and therapies that can target these potentially powerful effects are lacking.

In this study, we examined the effects of applied mechanical tension on breast cancer metastasis, growth and chemoresistance. We found that mechanical tension causes widespread changes in gene expression in breast cancer cells, leading to increased expression of genes relating to cell adhesion, drug metabolism and activation of Yap signaling and a reduction in genes for proliferation. On a functional level, mechanical tension increases the ability of breast cancer cells to adhere to specific extracellular matrix (ECM) molecules and TNF-α activated endothelial cells. Moreover, we found that mechanical forces could slow the proliferation of breast cancer cells and increase their resistance to a broad range of chemotherapeutic agents. We performed a screen of compounds to inhibit the activation of the cancer cells by mechanical load and found several inhibitors that were able to block mechanical force-induced enhancement of cancer cell adhesion under flow. Finally, we show that mechanically conditioned breast cancer cells have enhanced metastasis, altered growth and increased chemoresistance. In addition, we performed mechanobiological screening to find compounds that inhibited mechanical enhancement of metastasis and demonstrated that these compounds were active in a mouse model of metastasis. Together our findings support that mechanical tension is a powerful regulator of multiple aspects of breast cancer biology.

## Materials and Methods (Online)

### Cell Culture

Human umbilical vein endothelial cells (HUVECs) were cultured in MCDB-131 growth medium supplemented with 5% fetal bovine serum (FBS; Thermo Fisher Scientific, Inc.), SingleQuots growth supplements (Lonza), L-glutamine and penicillin/streptomycin. MDA-MB-231 and MCF-7 breast cancer cells were cultured in DMEM growth medium supplemented with 10% FBS, 1X MEM Non-Essential Amino Acids, L-glutamine and penicillin/streptomycin. MCF10A epithelial cells were cultured in DMEM/F12 medium, supplemented with 5% horse serum, 20 ng/mL EGF, 0.5 mg/mL hydrocortisone, 100 ng/mL cholera toxin, 10 µg/mL insulin, and penicillin-streptomycin. All cells were cultured at 37°C under a 5% CO_2_ atmosphere.

### High throughput cancer cell adhesion assay

In our past work, we developed a high throughput system for applying flow to cells cultured in 96 well plates (14–16). The device is composed of a rotational motor that drives a 96 gear box, simultaneously rotating 96 shafts with low angle cone tips, which interface with a standard format 96 well plate. As the low angle cones are brought into close proximity with the bottom of the wells, a linear shear stress is applied to the area of the well under the rotating cones. The high throughput cancer cell adhesion assay is performed by culturing HUVECs to confluence and then treating them with 10 ng/ml TNF-α for four hours prior to the adhesion assay. Cancer cells were labeled with CellTracker Dye (Thermo Scientific), trypsinized to detach them from the plate, and then allowed to recover in suspension for one hour prior to the start of the adhesion assay. We then added 1×10^5^ fluorescently labeled cancer cells/well to the well plate and applied 0.5 dynes/cm^2^ of shear stress for one hour. Non-adherent cells were washed from the well plate with PBS and remaining adherent cancer cells were detected using a plate reader. To assess the strength of adhesion of the adherent cancer cells, increasing bouts of shear stress were applied. Shear stresses of 1, 2, 3, 4, 5, 10 and 20 dynes/cm^2^ were applied for one minute increments, detached cells were washed from the plate, and the fluorescence of the remaining adherent cells were read by fluorescent plate reader after each level of shear stress.

### High throughput device for applying mechanical stretch to cultured cells

Our group has created a high throughput device that applies mechanical stretch to cancer cells in a 6 x 96 well format. Cells were cultured on custom 96-well culture plates with a flexible silicone membrane culture surface. Custom cell culture plates were assembled by sandwiching the silicone membrane between silicone gaskets, which were supported by polycarbonate wells and an aluminum base plate. The culture membrane was made of medical grade gloss/gloss 0.005” thick silicone. The well plates were sterilized by UV light and coated with collagen I (110 µg/mL) overnight. The well plates were mounted onto the stretch device and a true linear motor was used to drive platen with mounted pistons. Polytetrafluoroethylene pistons (5-mm diameter) on the platen were individually calibrated to apply precise amounts of equibiaxial strain to the silicone membrane. As the pistons are driven upward into the flexible membrane, the membrane is displaced a programmable distance which directly correlates to the application of a percent strain. The platen is mounted with linear ball bearings that run on six motion rails to guide precise alignment and motion of the platen. Vegetable oil was used to lubricate the pistons in contact with the silicone membrane of the culture plates.

### Adhesion and Transmigration Assays following Mechanical Loading

Silicone membrane bottom plates were coated with collagen I (110 µg/mL) overnight. Cancer cells were seeded at a concentration of 1×10^5^ cells/mL for 24 hours prior to application of strain. Cells were either cyclically strained at 0.1 or 1 Hz for 24 hours. Cancer cells were then trypsinized and allowed to recover for 1 hour. The suspension of cancer cells was added to a 96 well plate with confluent TNF-α activated HUVECs or purified extracellular matrix components fibronectin, collagen I, collagen II, collagen IV, vitronectin or laminin, and then adhesion and detachment assays were performed, as described above. The transmigration assay was performed using a Neuroprobe Chemotaxis Chamber. Endothelial cells were cultured to confluence on a porous membrane (8 µm diameter pores). Cancer cells were added to the top of the well in 5% serum MCDB 131 media, and 15% serum media was placed in the bottom of the well as a chemoattractant. MDA-MB-231 breast cancer cells were permitted to migrate for 16 hours and MCF-7 cells were permitted to migrate for 48 hours, then were fixed and assessed by plate reader and fluorescent imaging.

### Integrin and Kinase Inhibitor Library Treatment

MDA-MB-231 breast cancer cells were cultured on flexible membrane well plates, then treated with integrin inhibitors or kinase inhibitors for one hour before inducing cyclic mechanical strain. The integrin inhibitors used are listed in **Table 1**. The kinase inhibitor library (EMD Calbiochem; Cat. No. 539744) was used at a concentration of 10 µM for each inhibitor. After the addition of either integrin inhibitors or the kinase inhibitor library, cancer cells were mechanically strained for 24 hours. After mechanical strain, the cancer cells were detached from the well plate in trypsin for 2 minutes, spun down at 500g, resuspended and allowed to recover for 60 minutes in suspension. The cells were then used in the adhesion-detachment assays as described above.

### Immunostaining

After application of cyclic strain for 24 hours, the cells were washed with PBS and then fixed in 4% paraformaldehyde for 10 minutes. The cells were permeablilized in 0.2% Triton X-100 in PBS for 5 minutes and then blocked with 5% FBS in PBS and 1% BSA for 45 minutes. The cells were labeled with a 1:100 or 1:50 primary antibody dilution in PBS with 1% BSA overnight at 4°C. Cells were washed with PBS with 1% BSA, and labeled with fluorescent secondary antibodies and DAPI at a 1:1000 dilution (from a stock of 1 mg/ml) for 75 minutes at room temperature in the dark. The cells were washed extensively with PBS and mounted in mounting media (Vector Labs). The antibodies used in the studies are listed in **Supplemental Table 2**. For the studies with drug treatment and immunostaining the following inhibitors were used: Akt Inhibitor VIII (CAS 612847-09-3), JAK3 Inhibitor II (CAS 211555-04-3), JAK3 Inhibitor VI (CAS 856436-16-3), Lck Inhibitor (CAS 213743-31-8), PDGFR Tyrosine Kinase Inhibitor V (CAS 347155-76-4), N-Benzoyl Staurosporine (CAS 120685-11-2), PI3Kγ inhibitor (CAS 648450-29-7) and Verteporfin (CAS 129497-78-5).

### Western Blot Analysis

The cells were treated with the inhibitors described in the immunostaining experiments or on of the following inhibitors: Asciminib (CAS 1492952-76-7), Radotinib (CAS 926037-48-1), AZD1480 (CAS 935666-88-9), or Ruxolitinib (CAS 941678-49-5). Following treatments, the cells were washed with PBS and lysed using the following lysis buffer: 20 mM Tris (pH = 8.0), 5 mM EDTA, 150 mM NaCl, 1% Triton-X 100, 0.1% sodium dodecyl sulfate, 2 mM activated sodium orthovanadate, 1 mM phenylmethyl sulfonyl fluoride, 50 mM NaF, and a protease/phosphatase inhibitor cocktail (Thermo Scientific). Lysis buffer was added to the wells for 10 minutes, followed by scraping with cell scrapers. Lysate was alternately sonicated for 1 minute and kept on ice for 5 minutes for three cycles. The lysate was then centrifuged for 10 minutes at 10,000g. A protein assay was used to normalize the total protein in the samples (BCA assay; Thermo Scientific). The samples were mixed with 4X LDS Sample buffer in a 3:1 ratio with 5% β-mecaptoethanol. Samples were run on a precast NuPAGE Novex 4-12% Bis-Tris gels and transferred with iBlot transfer stacks to nitrocellulose membranes. The membranes were blocked in 5% milk in TBST or 5% StartingBlock T20 (Thermo Scientific) in TBST for one hour. The membranes were then incubated in 1° antibody in 1% milk or 1% StartingBlock T20 in TBST at 4°C overnight. Then the membrane was washed in TBST and incubated in 2° antibody (HRP conjugated) in 1% milk or 1% StartingBlock T20 in TBST. The antibodies used in the studies are listed in **Supplemental Table 3**. The membrane was washed extensively in TBST. The membranes were treated with luminol solution (SuperSignal West Femto Maximum Sensitivity Substrate; Thermo Fisher Scientific, Inc.) and then imaged using chemiluminescence imager (GBox-F3; Syngen Biotech.).

### RNA Sequencing and Analysis

MDA-MB-231 breast cancer cells were treated with mechanical loading for 24 hours and then RNA was isolated using the Qiagen RNeasy Mini Kit. The mRNA was sequenced using an Illumina HiSeq 4000. Single reads of 50 base pairs were performed after poly-A mRNA capture (Ambion Poly(A) Tailing Kit and NEBNext Ultra II Directional RNA Library Prep Kit) to isolate mRNA and dUTP directional preparation of the mRNA library. RNA sequencing was performed by the Genomic Sequencing and Analysis Facility at UT Austin. Gene expression analysis was performed using DESeq2 and R software. Plots were created using Prism GraphPad and Microsoft Excel. Gene ontology was performed using LAGO (Lewis-Sigler Institute, Princeton).

### Effect of Mechanical Strain on Chemotherapeutic Drug Response

MDA-MB-231 cells were plated on collagen I coated, custom made 96 well plates with flexible silicone bottoms. Cyclic mechanical strain at 7.5% or 15% was applied or cells were cultured statically for 8 hours. The cells were then dosed with paclitaxel, doxorubicin or 5-fluorouracil over a range of concentrations. Cells were strained for 16 hours followed by a second dose, and then an additional 24 hours of strain. At the conclusion of the assay, the cells were fixed with 4% paraformaldehyde, and the number of cells per field of view was quantified using DAPI staining.

### Multi Drug Resistance Flow Cytometry Assay

MDA-MB-231 breast cancer cells were mechanically strained at 0% and 7.5% strain for 24 hours. The EFLUXX-ID Gold multi drug resistance assay kit (Enzo Life Science) was used. For each sample, 5×10^5^ cells were treated with multi drug resistance inhibitors or untreated for 5 minutes, and then the Gold Efflux dye was added to all cells for 30 minutes. Propidium iodide was used to monitor cell viability. Cell fluorescence was then quantified via flow cytometry. Flow cytometry was performed using a LSRII Fortessa flow cytometer (BD Biosciences). FlowJo software (FlowJo, LLC) was used to analyze the results of the flow cytometry.

### Flow Cytometry Analysis of Cell Adhesion Markers

MDA-MB-231 breast cancer cells were mechanically strained at 0% and 7.5% cyclic mechanical strain for 24 hours. Following application of stretch, cancer cells were washed with PBS and detached from the flexible membrane using a cell scraper. The cells were centrifuged at 650g for 6 minutes and washed in cold BD Stain Buffer, then 1×10^6^ cells were labeled with fluorescently conjugated antibodies for 30 minutes at 4°C. The antibodies used are listed in **Supplemental Table 2**. Cells were washed twice in 1 mL of Stain Buffer. Cells were then fixed in 1% paraformaldehyde for 30 minutes at 4°C, then washed and stored in Stain Buffer until flow cytometry. Flow cytometry was performed using a LSRII Fortessa flow cytometer (BD Biosciences). FlowJo software (FlowJo, LLC) was used to analyze the results of the flow cytometry.

### Tube Formation Assay

Cancer cells were grown under static or mechanically loaded conditions for 24 hours. Cancer cells and HUVECs were labeled using CellTracker dye (Thermo Scientific) following the manufacturer’s instructions. HUVECs were seeded onto Matrigel in a glass bottom 96-well plate at a concentration of 20,000 cells per well. The HUVECs were treated with conditioned media from cancer cells by diluting the conditioned media in endothelial growth media in a 1:1 dilution. Cancer cells were seeded onto Matrigel at 20,000 cells per well into glass bottom 96-well plates. The formation of tubes in both assays were imaged using a Cytation 5 Cell Imaging Multi-Mode Reader (BioTek).

### Mouse xenograft model of metastasis

NOD/Scid mice (001303; Jackson Laboratory, Inc.) or nu/nu mice (002019; Jackson Laboratory, Inc.) were used in this study. For the xenograft study of breast cancer metastasis, MDA-MB-231 breast cancer cells were exposed to mechanical strain or static conditions for 24 hours. The cancer cells were either untreated or, treated with 10 μM of PI3-Kγ inhibitor (CAS 648450-29-7). We injected 5×10^5^ cells were injected into the tail vein of 7-week-old female mice. The mice were given an intraperitoneal injection of luciferin (150 mg/kg) in DPBS 10 minutes prior to imaging. The luminescence was visualized using the Xenogen IVIS Spectrum In Vivo Imaging System (PerkinElmer).

### Mouse orthotopic xenograft model

Nu/Nu mice (002019; Jackson Laboratory, Inc) were used in this study. For the xenograft study, luciferase-expressing MDA-MB-231 breast cancer cells were conditioned with mechanical strain or static conditions for 12 hours a day for 7 days. Following mechanical conditioning, 1×10^6^ cells were injected in 100 μl of Matrigel (Corning) into the inguinal mammary fat pad. Tumors were monitored daily and tumor growth was recorded with caliper measurements every other day. A laser speckle contrast imager (LSCI) was used in the orthotopic model to non-invasively quantify blood perfusion in the area of the injection. Imaging was performed as previously described with the entire back of the mouse captured in one image.(17, 18) The back of the mouse was illuminated with a diffuse laser diode (Thor Labs, 785nm, 50mW) to create a speckle pattern. The speckle pattern was captured using a Zoom-700 lens (Navitar) with a Bassler CCD (Graftek) and quantification was done using the contralateral side of the back as a relative control. To ensure comparability, both regions were captured simultaneously in one image.

### Micro-CT Analysis

The mice were imaged using micro-CT in the High Resolution X-Ray CT Facility at the University of Texas at Austin. The mice were perfusion fixed and then scanned using a micro-CT imaging system consisting of an NSI scanner, a Fein Focus High Power source, aluminum filter, and a Perkin Elmer detector. The scan was conducted at 140 kV, a source to object distance of 283 mm a source to detector distance of 1180 mm, and 0.25 pF gain. The scan was continuous with 2 frame averaging, 1800 projections, 5 gain calibrations, 5mm calibration phantoms and a 0.1 beam-hardening correction. Post reconstruction ring correction was applied using 2x over sampling, a radial bin width of 21 pixels, 32 sectors, a minimum arch length of 8 pixels, angular bin width of 9 pixels, angular screening factor of 4, and a voxel size of 51.5 µm. The raw CT slices were reconstructed using VGStudio 2.1, isolating the skull and preforming background subtraction. Light intensity and camera height were kept constant.

### Statistical Analysis

All results are shown as mean ± S.E.M., unless otherwise specified. Comparisons between multiple groups were analyzed using a two-way analysis of variance followed by a Tukey post hoc test. For studies that did not have normally distributed data, a Kruskal–Wallis test followed by multiple comparisons using the Conway-Imam procedure. Comparisons between only two groups with normally distributed data were analyzed using a Student’s *t* test. A two-tailed probability value *p* < 0.05 was considered statistically significant.

## Results

### Mechanical stretch dramatically alters gene expression profiles in breast cancer cells, including gene sets associated with Yap and Zeb transcription factors

We treated MDA-MB-231 cancer cells with mechanical strain (no strain, 7.5%, or 15% strain at 1 Hz) for 24 hours and then examined the change in gene expression using RNAseq. Mechanical loading at both magnitudes led to a broad shift in gene expression (**Fig. 1A, B; Supplemental Fig. 1**). Gene ontology of the significantly regulated genes identified significant upregulation in gene sets relating drug metabolism processes, angiogenesis, and regulation of proliferation and migration for cells treated with 7.5% strain (**Fig. 1C**). The most significantly altered gene sets were relating to fatty acid metabolism and other metabolism related gene sets for cells treated with 7.5% strain. At 15% strain, the most altered gene sets related to cellular differentiation, signaling and cell adhesion while the most downregulated genes were involved in morphogenesis, blood vessel development and epithelial branching (**Fig 1D**). Several genes associated with drug metabolism were also among the most upregulated genes in the entire genome, including Aldo-Keto Reductase Family 1 Member C1 (AKR1C1) that had a greater than 20-fold increase compared to static cultured cells (**Supplemental Fig. 1B**). We also observed large increases in related also-keto reductases AKR1B1 and AKR1B10. In addition, we found a decrease in expression of genes associated with proliferation compared to statically cultured cells (**Supplemental Fig. 1C**). Many genes known to be controlled by Yap and Zeb transcription factors were also significantly regulated at both 7.5% and 15% strain compared to static cells (**Supplemental Fig. 1D**). In addition, there was a significant shift in expression for genes relating to cell-cell adhesion and cell adhesion to extracellular matrix (**Supplemental Fig. 1E**).

**Figure 1.**
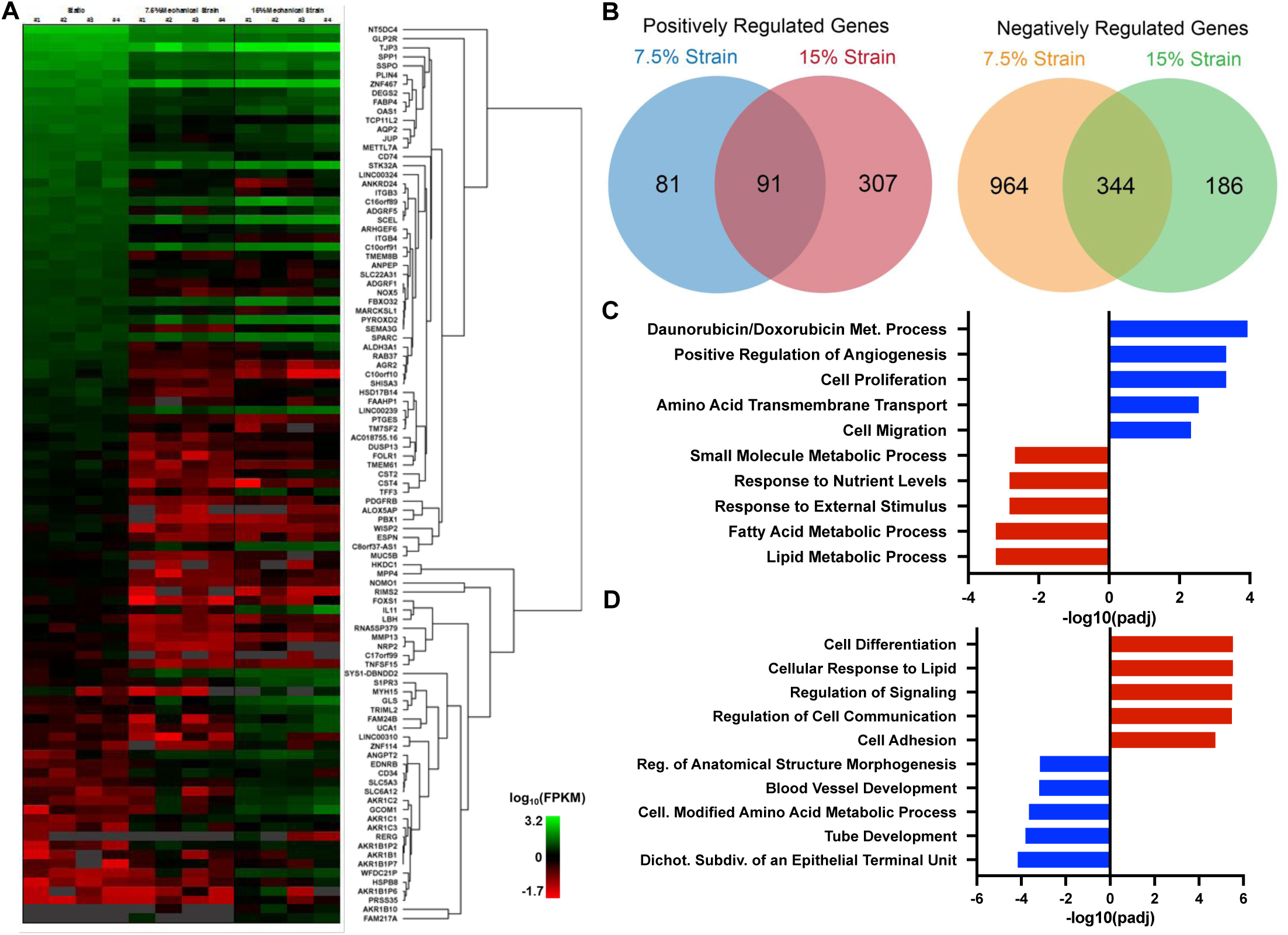
Mechanical strain regulates gene transcription of cell adhesion, drug metabolism and proliferation genes. MDA-MB-231 breast cancer cells were treated with mechanical strain for 24 hours at 0, 7.5, and 15% maximal strain at a frequency of 1 Hz. Total RNA was isolated and RNA sequencing was performed (n = 4). (A) Hierarchical clustering of the most significantly regulated genes. FPKM = fragments per kilobase of transcript per million mapped reads. (B) Venn diagrams for significantly regulated genes in the 7.5 % and 15% mechanical strain groups. (C) The top five most upregulated and downregulated gene ontology groups for cells treated with 7.5% strain. (D) The top five most upregulated and downregulated gene ontology groups for cells treated with 15% strain.

### Mechanical strain decreases proliferation and apoptosis signaling and increases breast cancer resistance to chemotherapeutic drugs

Using a high throughput mechanical loading system recently developed by our laboratory (19, 20), we conditioned cancer cells with a physiological range of mechanical strains ranging from 0 to 17.5% strain to MDA-MB-231 breast cancer cells. We found that mechanical loading at any of the magnitudes reduced expression of the proliferation marker Ki-67 (**Fig. 2A, B**). Overall metabolic activity as measured by an MTT assay was over 50% lower in cells under 7.5% mechanical strain in comparison to cells grown under static conditions (**Supplemental Fig. 2**). Immunostaining for anti-apoptotic Bcl-2 and pro-apoptotic Bax showed a reduction in the ratio between Bax and Bcl-2 with mid-range mechanical strains (**Fig. 2C, D**). Gene expression for Bax and Bcl-2 RNAseq was consistent with these findings, showing increases in both the Bax and BCL2 genes however there was no alteration in the ratio of Bax to BCL2 in gene expression (**Fig. 2E**). To examine, the effect of extracellular forces can alter chemoresistance, we applied chemotherapeutic compounds to the cells in combination with 7.5% or 15% cyclic mechanical strain. We found mechanically loaded cells had reduced proliferation however there was little alteration in the proliferation of loaded cells even in the presence of high doses of paclitaxel, doxorubicin and 5-fluorouracil (**Fig. 2F**). A multidrug resistance assay showed that mechanical conditioning at 7.5% strain did not affect MDR1, MRP1/2 and BCRP drug efflux pump activity (**Supplemental Fig. 3**). In previous studies, alterations in the Abl1 signaling in breast cancer cells led to increased chemoresistance while reducing proliferation (21, 22). Abl1 is also associated with the cytoskeleton and is mechanically regulated in other cell types (23, 24). We applied 7.5% mechanical strain to the cells with or without inhibitors to Abl1, Jak2 or PI3K and then performed immunoblotting to cell survival and signaling pathways (**Fig. 2G, H; Supplemental Fig. 4**). Mechanical loading led to increases in phosphorylated Bcl-xL, p-p70/85 S6K, Akt, Jak2, PI3K, and Mcl1. In addition, there were increases in total p70 S6K and a reduction in cyclin D1. Inhibitors to Abl1 reduced the load-induced phosphorylation of Bcl-xL, PI3K and Mcl1 while inhibitors to Jak2 or PI3K reduced load-induced phosphorylation of Akt (**Fig. 2G, H; Supplemental Fig. 4**). Together, these findings suggest a load activated signaling pathway in which Abl1 and Jak2 are activated by mechanical load to induce increases in pro-survival signaling (**Supplemental Fig. 5**).

**Figure 2.**
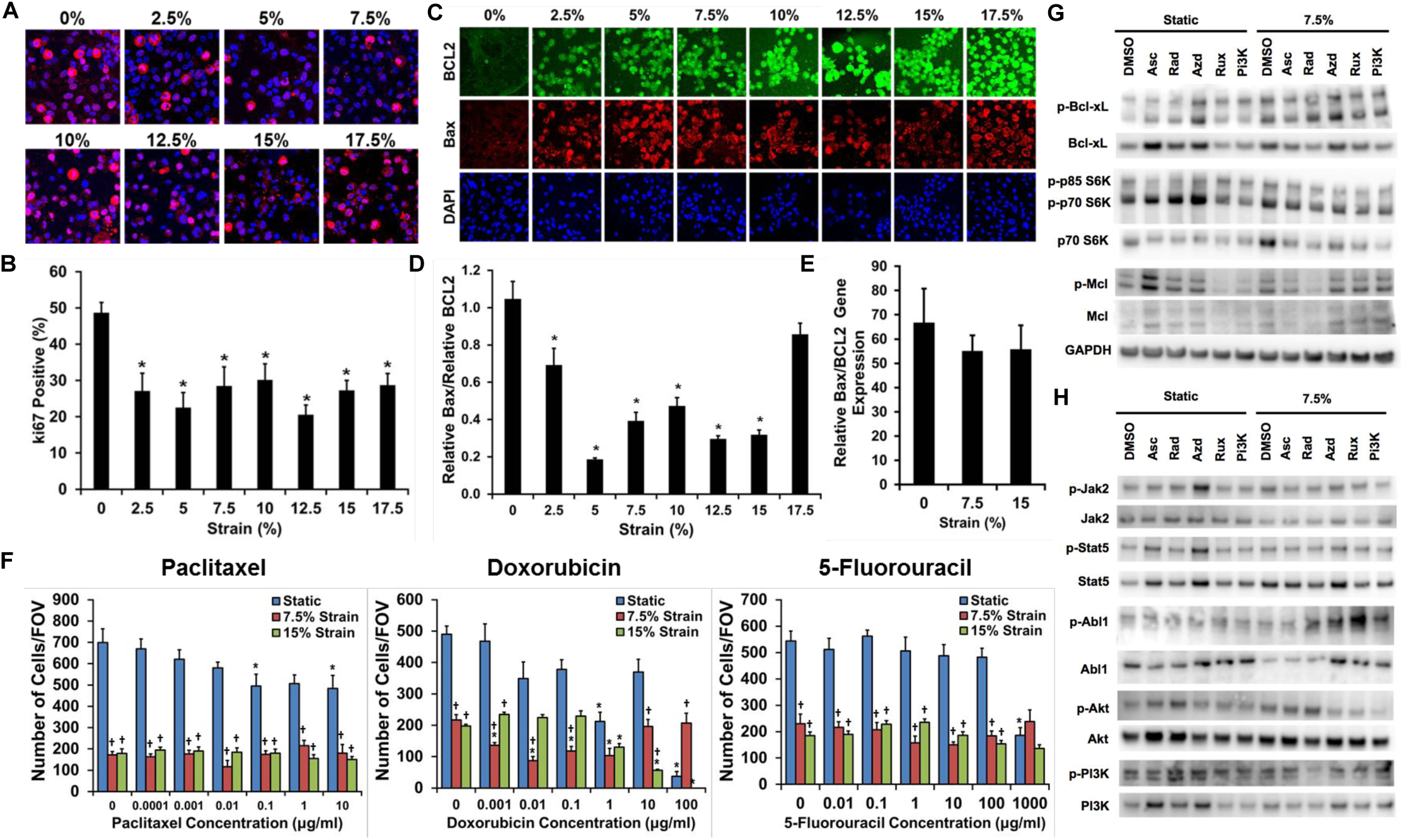
Mechanical strain decreases proliferation and increases drug resistance in breast cancer cells. (A) MDA-MB-231 breast cancer cells were mechanically strained at 1 Hz for strains ranging from 0 to 17.5% for 24 hours and then immunostained for Ki-67. Bar = 100 μm. (B) Quantification of Ki-67 positive cells following mechanical loading. *p* < 0.05 versus static culture (0% strain; n = 8). (C) Images of Bax and Bcl-2 immunostaining for cyclic mechanical strains of 0 to 17.5% strain. Bar = 100 μm. (D) Relative expression of pro-apoptotic Bax to anti-apoptotic Bcl-2 protein expression. *p < 0.05 versus 0% strain (n = 10). (E) Relative gene expression of Bax to Bcl-2 after 24 hours of mechanical strain (n = 10). (F) Response of MDA-MB-231 breast cancer cells to drug treatment with paclitaxel, doxorubicin, or 5-fluorouracil under 0%, 7.5%, or 15% mechanical strain (n = 8). *p < 0.05 versus static conditions. ^†^p < 0.05 versus static conditions under no treatment and under static conditions with the pharmacological treatment with same concentration as the indicated group. (G) Western blotting for cells treated with 7.5% strain in combination with DMSO, asciminib (Asc; Abl1 inhibitor), radotinib (Rad; c-Abl1 inhibitor), AZD1480 (Azd; Jak2 inhibitor), Ruxolitinib (Rux; Jak1/2 inhibitor), or a PI3K inhibitor (PI3K).

**Figure 3.**
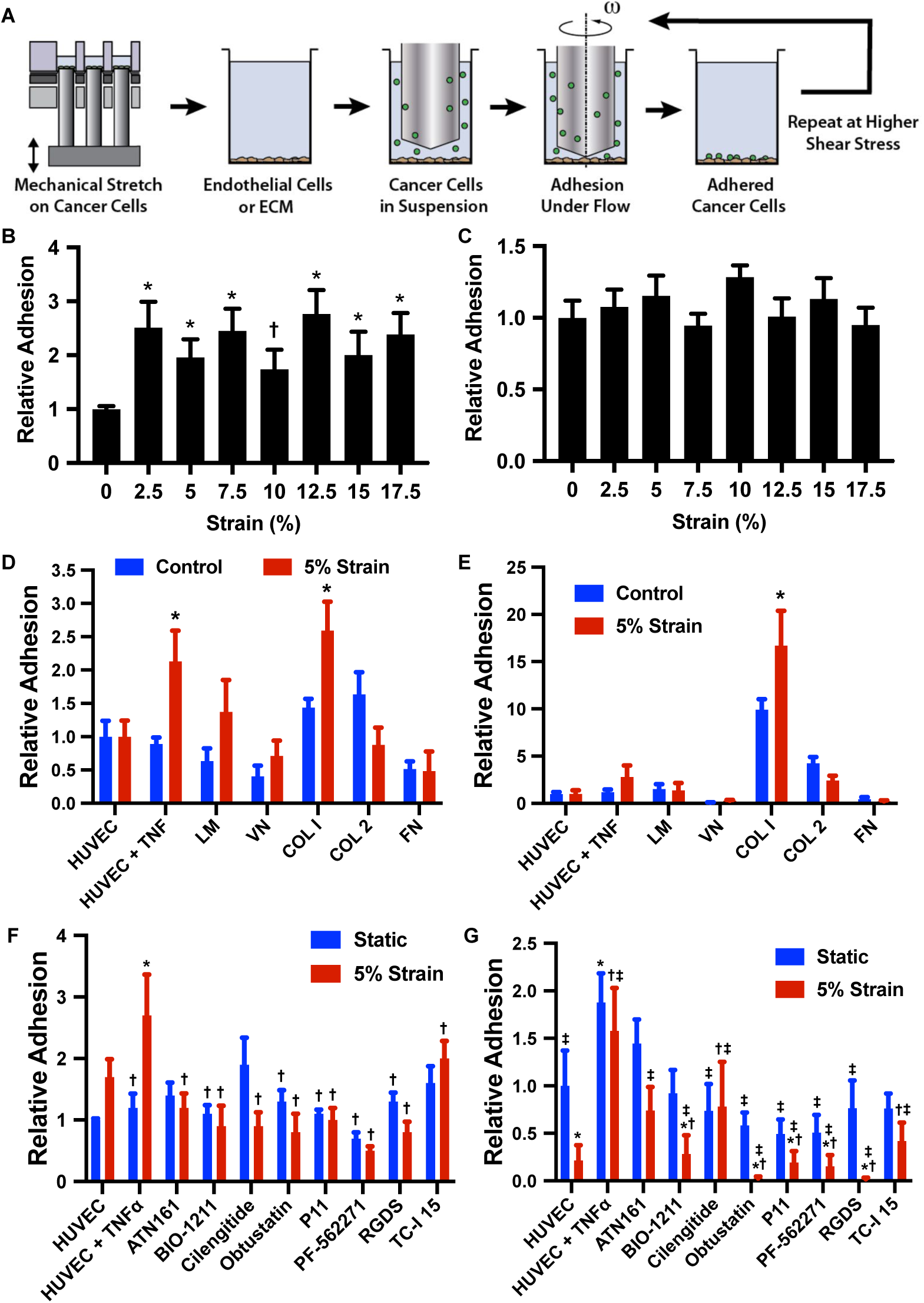
Mechanical load enhances the adhesion of cancer cells to endothelial cells and collagen I under shear stress. (A) Diagram of the experimental design. The cells were first treated with mechanical load in a high throughput system and then adhesion measured in a high throughput flow device. (B) MDA-MB-231 and MCF-7 breast cancer cells were mechanically strained at maximal strain from 0% to 17.5% at a frequency of 0.1 and 1 Hz for 24 hours. Initial adhesion of strained cells under 0.5 dynes/cm^2^ shear stress to a TNF-α treated endothelial monolayer was measured relative to the static group. \**p* < 0.05 compared to the static group (n = 8). (C) Relative adhesion of the cells treated with mechanical load after detachment shear stress up to 20 dynes/cm^2^. (D) Adhesion of cells to endothelial cells and isolated ECM molecules including laminin (LM), vitronectin (VN), collagen I (COL I), collagen II (COL 2) and fibronectin (FN). *p < 0.05 versus control group with the same adhesion substrate. (E) Adhesion of cells to endothelial cells and ECM after detachment with shear stresses up 20 dynes/cm^2^ (n = 8). *p < 0.05 versus control group with the same adhesion substrate. (F) Initial adhesion of cells to endothelial cells in the presence of integrin inhibitors. Adhesion is to endothelial cells treated with TNF-α unless otherwise noted. (G) Relative adhesion of cells after detachment up to 20 dynes/cm^2^ (n = 8). Adhesion is to endothelial cells treated with TNF-α unless otherwise noted.*p < 0.05 versus the HUVEC group. ^†^p < 0.05 versus versus the TNF-α treated HUVEC group.^‡^p < 0.05 versus the TNF-a treated HUVEC with 5% strain group.

**Figure 4.**
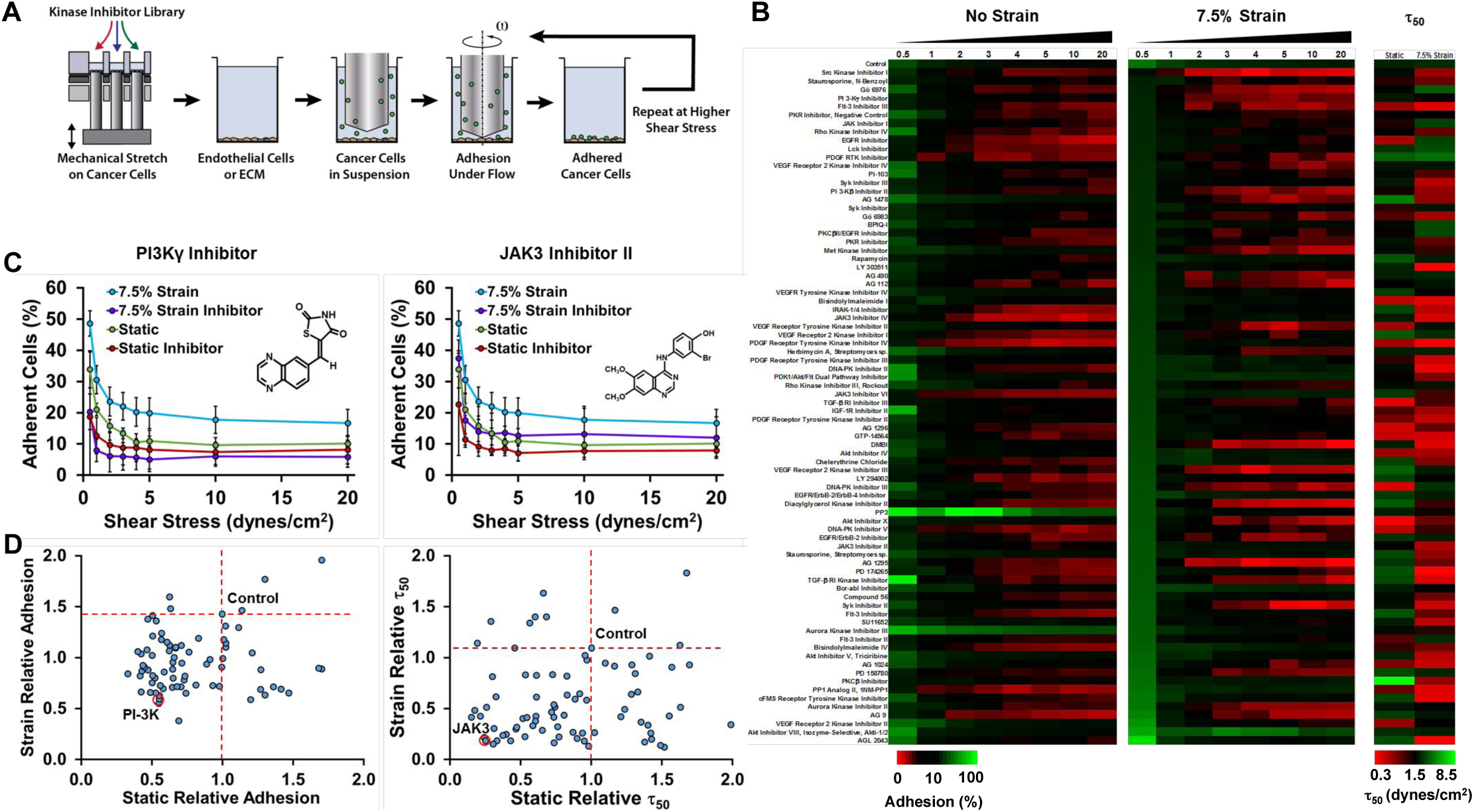
High throughput mechanobiological screens for blocking load-induced enhancement of cancer cells to endothelial cells under shear stress. (A) Diagram of experimental protocol. The cells are loaded in the presence of compounds from a kinase inhibitor library and then the adhesion to inflamed endothelial cells under flow is measured. (B) Heat map of the adhesion and detachment of cancer cells with treatment with kinase inhibitors. (C) Example graphs from the kinase screen for a PI3K inhibitor and Jak3 inhibitor. (D) Results of the kinase inhibitor screen. Compounds in the lower left portion of the graph have reduced initial adhesion or τ_50_ to endothelial cells. The τ_50_ is an index of the strength of adhesion. It is the shear stress needed to cause detachment of 50% of the cells calculated from a curve fit to the detachment of the cells under increases shear stress.

**Figure 5.**
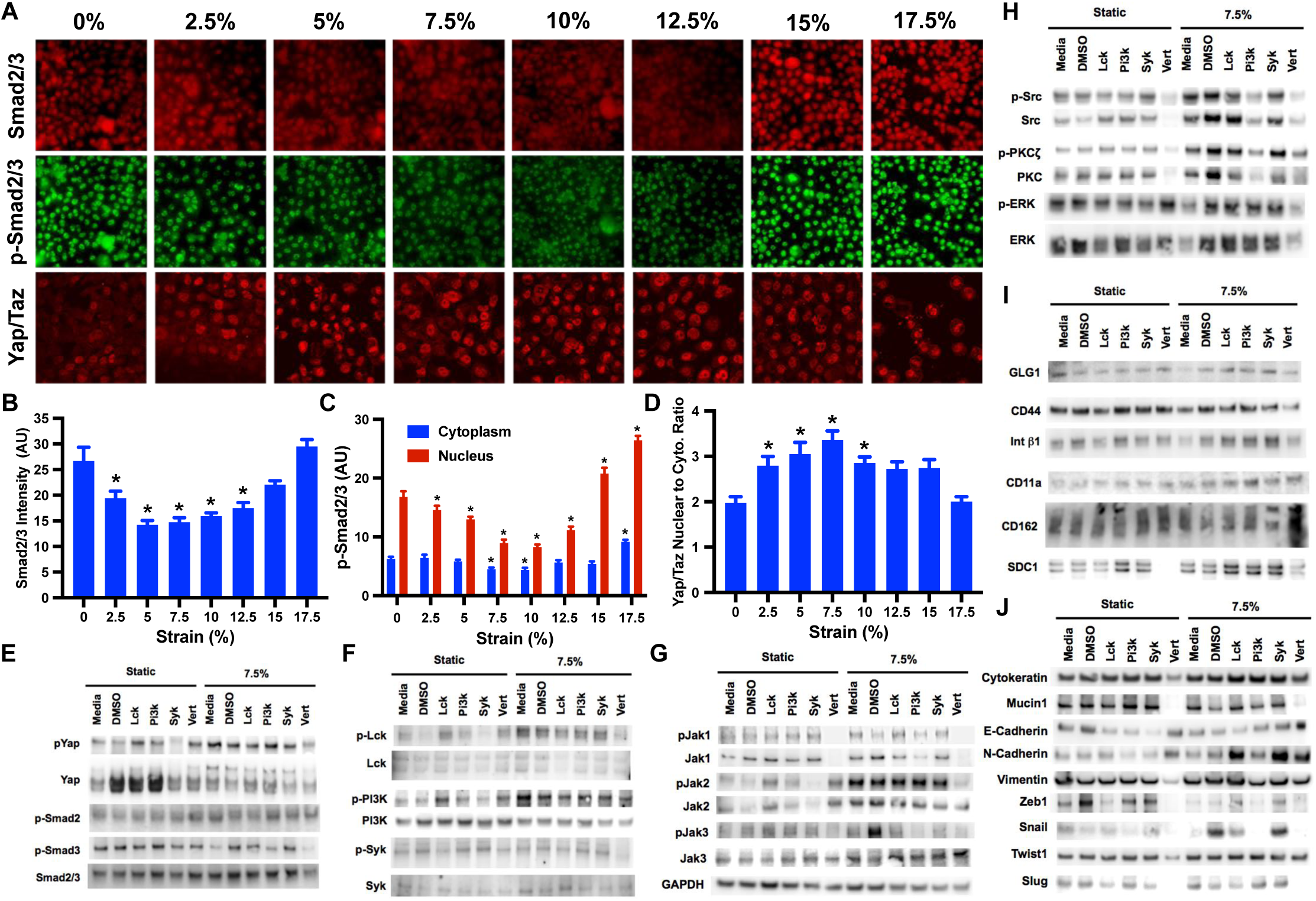
Strain differentially activates Smad2/3 and Yap/Taz in breast cancer and epithelial cells. (A) MDA-MB-231 cells were treated with mechanical strain for 24 hours at 1 Hz with varying magnitudes of maximal strain. Immunostaining for phospho-Smad2/3, Smad2/3 and Yap/Taz was performed. Bar = 100 μm. (B) Quantification of total Smad2/3 in the cells (n = 20). \**p* < 0.05 versus static group (n = 20). (C) Quantification of phosphorylated Smad2/3 in cells treated mechanical load (n = 20). *p < 0.05 versus the static group for the same subcellular location. (D) Quantification of nuclear to cytoplasmic Yap/Taz staining (n = 20). \**p* < 0.05 versus static group. (E-J) Western blotting for lysates from cells treated with mechanical load for 24 hours and inhibitors Lck, PI3K, Syk and Yap (Vert; verteporfin).

### Mechanical strain increases adhesion of breast cancer cells to endothelial cells and collagen I

A key step during metastasis is the adhesion and arrest of circulating cancer cells to the endothelium or subendothelial matrix (25, 26). Using the high throughput mechanical loading system, we applied mechanical stretch to MDA-MB-231 breast cancer cells over a range from 0 to 17.5% maximal strain for 24 hours. Our group has developed a high throughput flow system that allows the application of controlled flow in a 96 well plate format (14,16,27). The system uses low angle cones rotated near the culture surface of the well to produce shear on the cells. We performed assays for cell adhesion under flow using the high throughput flow system after applying mechanical load (**Fig. 3A**). After initial adhesion under flow, we washed the plates and then applied progressively increasing shear stress in short bouts to detach the cells and assess the strength of adhesion. We found that all levels of strain tested increased adhesion to endothelial cells pretreated with TNF-α (**Fig. 3B**). There were no significant changes in the strength of adhesion for the strained cells (**Fig. 3C**). To test the specificity of adhesion induced by mechanical loading we repeated the experiment exposing the cells to 5% strain for 24 hours and then tested adhesion to non-TNF-α treated endothelial cells and purified ECM molecules. We found that there was increased initial adhesion to TNF-α treated endothelial cells and to collagen I but not to other ECM molecules or non-inflamed endothelial cells (**Fig. 3D**). There was an increase in the strength of adhesion of cells to collagen I but not to other ECM molecules or endothelial cells (**Fig. 3E**). We next repeated the studies and included the treatment with integrin and focal adhesion kinase (FAK) inhibitors during the mechanical strain (but not during the adhesion assay). Inhibitors to β1 integrins, αvβ3/β5 integrins, pan-integrin inhibitor and the FAK inhibitor, reduced the mechanical strain strain-enhanced cell adhesion and adhesion strength to endothelial cells (**Fig. 3F, G**).

### High throughput mechanobiological screen identifies kinase inhibitors that inhibit mechanical strain enhanced cancer cell adhesion to inflamed endothelial cells

We next performed a high throughput screen to identify potential pathways and compounds that could be used to inhibit the load-induced enhancement of cancer cell to endothelial cell adhesion under flow. We treated cancer cells with mechanical load at 7.5% strain at 1 Hz under treatment with compounds from a drug library of kinase inhibitors. After 24 hours, we performed a cell adhesion assay under shear stress using the high throughput flow system to identify compounds that decreased cancer cell adhesion under both static and mechanical loading conditions (**Fig. 4A**). The screen allowed an assessment of initial adhesion as well as an index for strength of adhesion by calculating a shear stress at which 50% of the cells would be removed (τ_50_; **Fig. 4B-D**). Inhibitors for EGFR reduced adhesion of cells cultured statically (10 of 12 inhibitors) and under mechanical load (5 of 12 inhibitors). Under static conditions, inhibitors for FLT-3 (3 of 3 inhibitors) and PDGFR (6 of 9 inhibitors) reduced adhesion but not for cells treated with mechanical load. Inhibitors for JAK3 (2 of 3 inhibitors), PI3K (2 of 4 inhibitors), Syk (2 of 3 inhibitors) and VEGFR (3 of 6 inhibitors) reduced adhesion to varying degrees in cells cultured under static or mechanical loading conditions. For inhibiting adhesion strength, inhibitors of AKT (3 of 3 inhibitors), EGFR (9 of 12 inhibitors), JAK3 (3 of 3 inhibitors), PKC (3 of 4 inhibitors) and Rho kinase (2 of 2 inhibitors) were effective in blocking adhesion strength of loaded and non-loaded cells. Two JAK3 and two Rho kinase inhibitors were among the inhibitors that most decreased the τ_50_. In general, for the library, the inhibitors studied were more effective at reducing adhesion under the static conditions in comparison to loading conditions. Out of a total of 80 inhibitors, 21 inhibited adhesion of statically cultured cancer cells by at least 45% whereas only 4 inhibitors were able to reduce adhesion on loaded cells by the same amount.

### Mechanical strain increases production of angiogenic soluble factors and endothelial cell-like behavior in cancer cells

The gene expression analysis had revealed increases in angiogenesis-related genes in the cancer cells including endothelin-1 (EDN1), follistatin (FST), GATA-binding protein 2 (GATA2), Kruppel like factor 5 (KLF5), Platelet-Derived Growth Factor A (PDGFA), tumor necrosis factor receptor superfamily member 12A (TNFRS12A), and Rho GTPase Activating Protein 22 (ARHGAP22; **Supplemental Fig. 1**). To test the effect of mechanical loading on the angiogenic properties, we exposed MDA-MB-231 cells to mechanical load for 24 hours (1 Hz at 7.5% strain) and then assayed their angiogenic properties in a tube formation assay. The conditioned media from loaded cells stimulated increased tube formation in endothelial cells (**Supplemental Fig. 6A**). In addition, cancer cells formed more tubes when seeded on matrigel with mechanical conditioning (**Supplemental Fig. 6B**), indicative of increased potential for vascular mimicry for the loaded cells (28).

**Figure 6.**
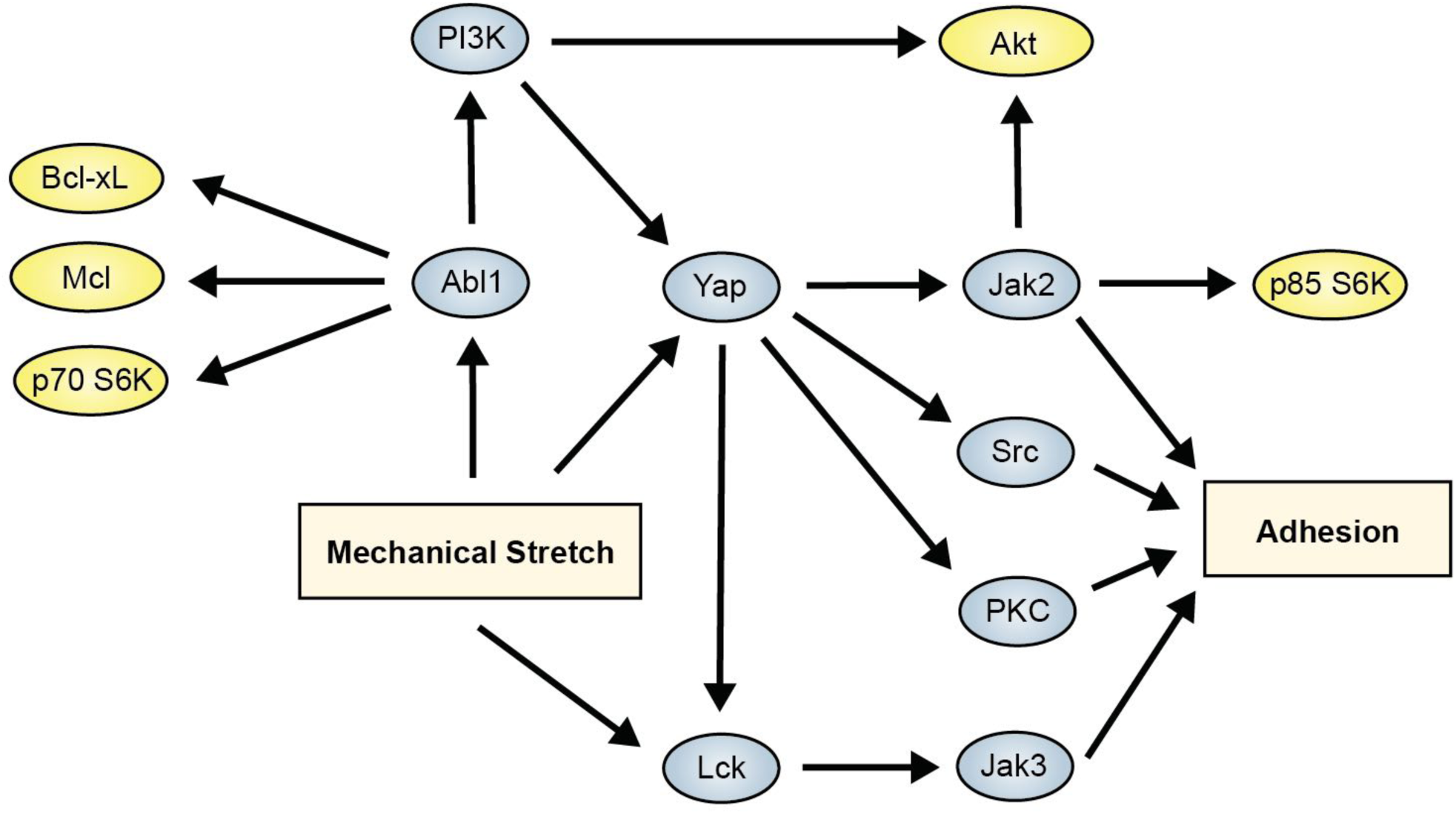
Summary diagram of the mechantransduction mechanisms supported by the studies for the enhancement of survival and adhesion by mechanical load.

### Mechanical strain increases adhesion through Abl1, Yap and Lck-mediated pathways

To understand the mechanotransduction pathways that were key in regulating load-induced increases in cancer cell adhesion under flow, we examined the activation of adhesion related pathways that were implicated by our studies on cancer cell survival and pathways that were indicated from the “hits” from the kinase inhibitor screen. We applied mechanical load to the MDA-MB-231 cells at varying magnitudes for 24 hours (1 Hz). Immunostaining indicated that there was a decrease in phosphorylated Smad2/3 at mid-level strains (**Fig. 5A-C**). In addition, at mid-level strains there was also a significant increase in the nuclear cytoplasmic ration of Yap/Taz (**Fig. 5A, D**). Western blotting to lysates from the loaded cells showed increases in phospho-Yap, phospho-Smad3 and an increase in phosphorylation for phospho-Smad2 (**Fig. 5E**).

In addition, mechanical load led to activation of Lck, Jak1-3, Src, and PKC pathways (**Fig. 5F-5H**). Using inhibitors to Lck, PI3K, Syk and Yap, we demonstrated that many of the pathways were dependent on the activation of Yap (**Fig. 5E-H**). Yap is key to many of the adhesion-mediated pathways but Lck signaling provides a Yap-independent mechanism for enhancing load-induced adhesion. This pathway is also consistent with our survival studies as Abl1 can be induced by load to increase pro-survival signaling even in the presence of Yap or Jak2 inhibition. We examined the levels of several adhesion receptors that could be involved in enhanced endothelial cell adhesion under mechanical load, however, among these only syndecan-1 (SDC1) was upregulated and a decrease in expression did not correlate with the decreased expression from the inhibitor studies (**Fig. 5I**). We also examine molecules and signaling related to epithelial-to-mesenchymal transition (EMT). While there were changes with load in some of the molecules, there was not a clear shift to epithelial or mesenchymal phenotype with mechanical loading (**Fig. 5J**). Combined with our results in the studies on cell survival, these results a signaling cascade in which mechanical load activates multiple pathways that enhance both survival and adhesion (**Fig. 6**).

### Mechanical strain induces increased metastasis to the skull, angiogenesis and reduces tumor growth in mouse xenograft models

We next examined whether mechanical conditioning of cancer cells led to altered behavior in *in vivo* models of tumor growth and metastasis. We mechanically conditioned MDA-MB-231 cells with 7.5% strain at 1 Hz for 24 hours and then implanted them in an orthotopic xenograft tumor model in the mammary fat pad of nu/nu mice. We found that there was decreased tumor growth by IVIS imaging for the cells that had been mechanically conditioned in comparison to those grown under static conditions (**Fig. 7A, B**). The tumor volume measured by calipers was significantly lower in the mechanically conditioned cells in comparison to the controls on day 3; however, there was not a significant difference between static and mechanically loaded cells at other time points (**Supplemental Fig. 7A, B**). There were also no significant differences in the weight of the tumors on explantation for the comparison between static or mechanically loaded cells at the final time point (**Supplemental Fig. 8**). Using laser speckle imaging we examined the relative perfusion between the mammary fat pads in the mice. We found significant increases in perfusion in the fat pads of mice with mechanically loaded cancer cells at day 2 and day 8 following implantation (**Fig. 7C, D**). At later time points, there was no difference in the perfusion between the fat pad with static or mechanically loaded cancer cells (**Supplemental Fig. 9**). To test whether mechanical conditioning predisposed cancer cells to metastasize, we mechanically conditioned the cells for 24 hours and then injected them through the tail vein in nu/nu or NOD Scid mice. In both the models, we did not observe a significant increase in lung metastases. However, in both models, there was increased tumor colonization of the skull (**Fig. 7E-H**). When cells were pretreated with a PI3Kγ inhibitor identified from our high throughput screen we found decreased metastasis in the mechanically loaded group **(Fig. 7G, H**). MicroCT analysis of the skull revealed small regions of bone loss in the mice injected with mechanically conditions cells in the occipital and postorbital bones (**Supplemental Fig. 10**).

**Figure 7.**
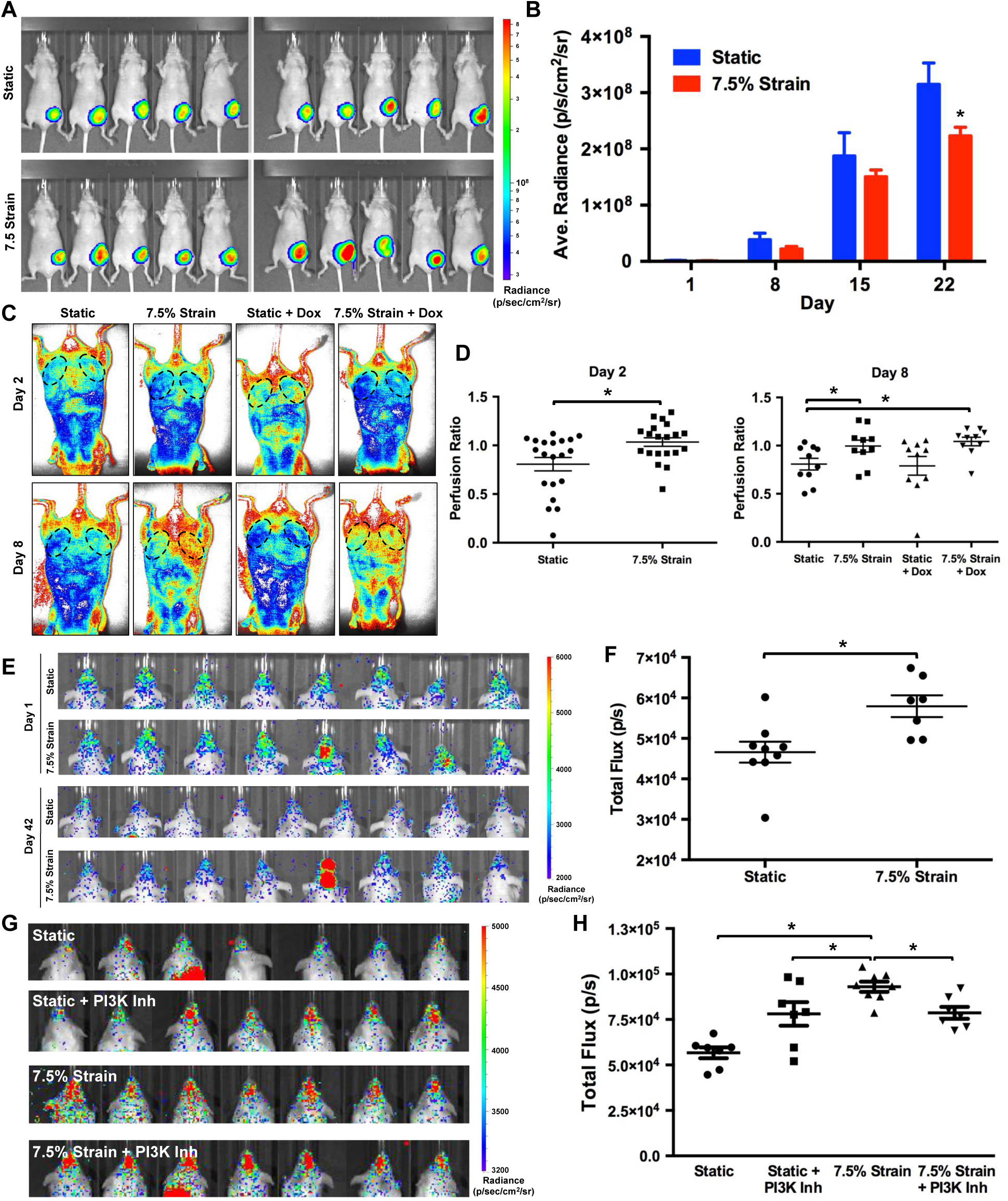
Mechanical loading decreases tumor growth and increases metastasis to the skull in immune compromised mice. (A) Luciferase expressing MDA-MB-231 cells were grown under static or mechanically loaded conditions for 24 hours and then implanted into the mammary fat pad of nu/nu mice. Radiance of mice with orthotopic tumor implantation at 22 days. (B) Quantification of the radiance in the mammary fat pad. *p < 0.05 versus static group (n = 10). (C) Laser speckle imaging of mice with orthotopic implantation of MDA-MB-231 cells after mechanical conditioning. (D) Ratio of perfusion of the mammary fat pad with tumor implantation to contralateral control fat pad. *p < 0.05 between indicated groups (n = 10). (E) Visualization of luminescence by IVIS for nu/nu mice given a tail vein injection of MDA-MB-231 cells cultured under static or 7.5% strain for 24 hours. (F) Quantification of luminescence in the head of the mice at day 42 following injection. *p < 0.05 versus static group (n = 8-9). (G) Luminescence in the heads of NOD Scid mice 42 days after tail vein injection of MDA-MB-231 cells. (H) Quantification of the luminescence in the head of the mice after 42 days. *p < 0.05 versus indicated group (n = 7-8).

## Discussion

While many studies have supported the role of matrix stiffness in regulating cancer biology (4,29–31), the role of applied forces on tumor biology is far less well understood. Physical activity has been linked to improved outcomes in breast cancer by many studies. Recent meta-analyses concluded that regular exercise reduces the mortality from breast cancer by 40% and that physical activity was the most powerful lifestyle-based modifier for breast cancer outcomes (32, 33). While the benefits of exercise in cancer patients would certainly be multifactorial, our study sought to examine whether forces applied to breast cancer cells could directly alter their survival and metastasis. Our findings support that applied cyclic tension induces pro-survival signaling through the Abl1 and Jak2 pathways while reducing proliferation of the cells. This pro-survival signaling enables the tumor cells to survive with level of chemotherapeutics that would otherwise cause cell death. In addition, through a set of mechanobiological screens we identified pathways that underlie load-induced enhancement of cancer cell adhesion to endothelial cells under flow. These pathways included those that were both dependent and independent of Yap signaling.

A major finding of our study is the development of chemoresistance in the mechanically loaded cells through multiple pathways. Notably, in our gene expression analysis several of the aldo-keto reductase family members, related to drug metabolism, were among the most upregulated genes by mechanical load out of the entire genome. In addition to the pro-survival signaling that was increased by mechanical load, upregulation of drug metabolism would further provide a mechanism for chemoresistance. Parallel to our findings, another study found that A549 lung cancer cells when mechanically strained at 20% maximal strain for 6 days, had reduced proliferation and responsiveness to chemotherapeutic drugs (34). However, in another study in hepatocellular liver carcinoma cells in which the cells were exposed to shear stress from orbital shaking, there was increased cell death with cisplatin treatment in combination with shear stress (35). In MDA-MB-231 cells, 15 minutes of vibrational stresses per day also reduced proliferation (12). While compressive stress was found to enhance invasion in several cancer cell lines including the MDA-MB-231 cell line (8). In addition, compressive forces can suppress proliferation and induce apoptosis in cancer cells (13). Our *in vivo* study did not show differences in tumor growth under doxorubicin treatment. However, our study had the limitation that the loading was only applied prior to implantation and thus this would suggest that effects of load are reduced over time. This concept is further supported by our finding that angiogenesis was increased in loaded cells at early time points but not for later time points. From these studies, one could infer the timeline of the effects of load last around one week. A recent study found that mice that performed stretching exercises after orthotopic implantation of breast cancer cells also had reduced tumor growth (36). These findings are consistent with our results and support that breast cancer growth can be reduced by mechanical stretch.

Our studies found enhanced metastasis to the skull of mice injected with mechanically loaded cells. Yap/Taz pathway activation has been observed in high-grade breast cancer metastatic cancer in comparison to non-metastatic cancer (37). Taz is also required for metastasis in breast cancer stem cells (38). Activation of Yap pathway target genes induces bone metastasis in breast cancer through signaling pathways involving ROR1, HER3 and lncRNA MAYA (39). In our studies, we found Yap-dependent activation of Src, PKC and Jak2 was partially responsible for load induced increases in adhesion. However, there appeared to be a separate mechanism acting through Lck and Jak3 that also increased adhesion following loading. Thus, these findings are consistent with previous work suggesting Yap activation increases bone metastasis but also suggests that targeting Yap would not be sufficient to completely block this effect. Our screen for metastasis inhibitors identified inhibitors to several pathways that reduced the adhesion of cancer cells with and without mechanical preconditioning. These included inhibitors to FLT-3, PI-3K, JAK3, EGFR and VEGFR. Consistent with our findings here, our previous study using the high throughput shear stress device also identified FLT-3 inhibitors to be effective in blocking cancer cell adhesion to endothelial cells under shear stress (15). The PI3K inhibitor reduced metastasis to the skull in immune compromised mice for the load conditioned cells, suggesting the strategy of screening for inhibitors of cancer cell to endothelial cell adhesion could be a means to identify therapeutics that protect against metastatic spread of cancer but that multiple pathway targeting may be needed.

Overall, our studies support the concept that microenvironmental mechanical stresses can regulate many aspects of breast cancer biology. The mechanical loading environment of tumors in the body is highly variable and there is potential that tumor cells that are in locations with specific mechanical forces may be predisposed to survive during chemotherapy, to metastasize or induce the development of vasculature. Thus, there is a need to consider the effects of mechanical forces on tumor behavior when designing and testing therapeutics for cancer or developing accurate models of tumor growth.

## Acknowledgements

The authors gratefully acknowledge funding through the American Heart Association (17IRG33410888), the DOD CDMRP (W81XWH-16-1-0580; W81XWH-16-1-0582) and the National Institutes of Health (1R21EB023551-01; 1R21EB024147-01A1; 1R01HL141761-01) to ABB. The authors also acknowledge support through a graduate fellowship from the NSF (DGE-1610403) to A.S.

## Supplemental Figure Legends

**Supplemental Figure 1.**
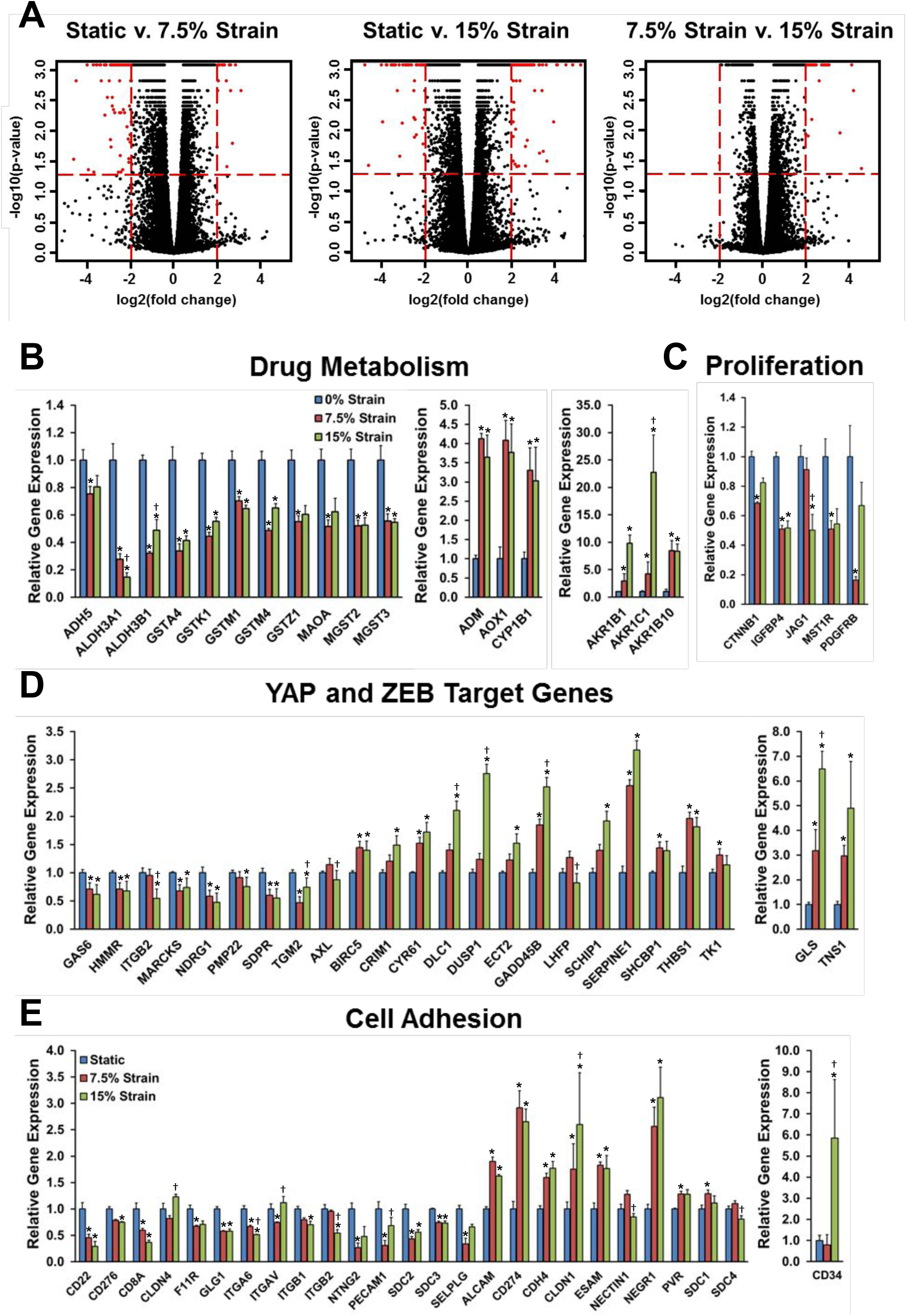
Mechanical strain regulates gene expression in functional gene groups. MDA-MB-231 breast cancer cells were mechanically strained at 0, 7.5 or 15% strain at a frequency of 1 Hz for 24 hours (n = 4). (A) Volcano plots for comparisons between the groups. (B-E) RNA sequencing was performed and gene groups were examined including genes associated with drug metabolism, proliferation, Yap/Zeb transcriptional control, and cell adhesion. *p < 0.05 versus static group. **^†^**p < 0.05 versus 7.5% strain group.

**Supplemental Figure 2.**
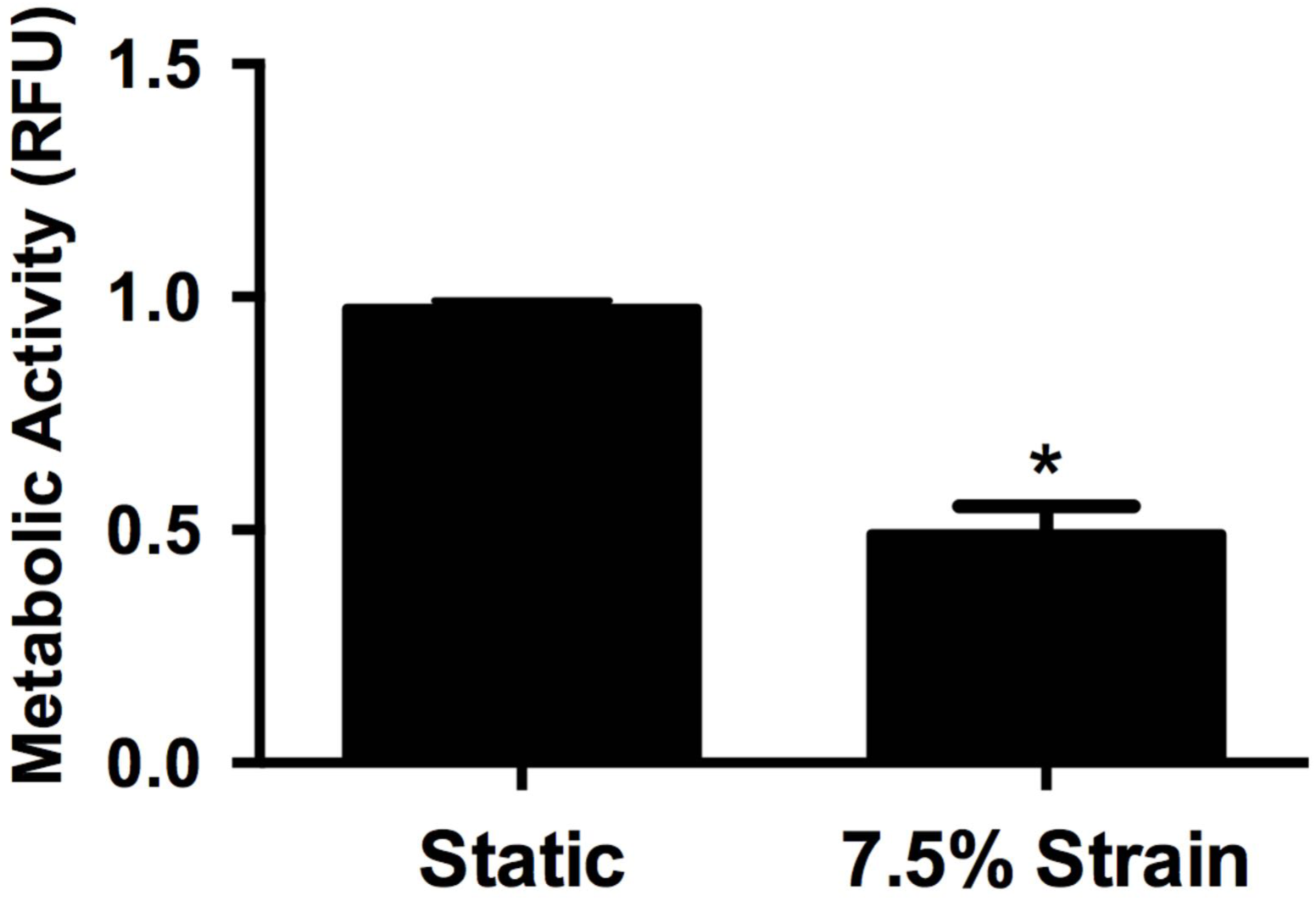
MDA-MB-231 cells were treated with 7.5% mechanical strain for 24 hours and metabolic activity was measured using an MTT assay. \**p* < 0.05 versus 0% strain group (n = 6-8).

**Supplemental Figure 3.**
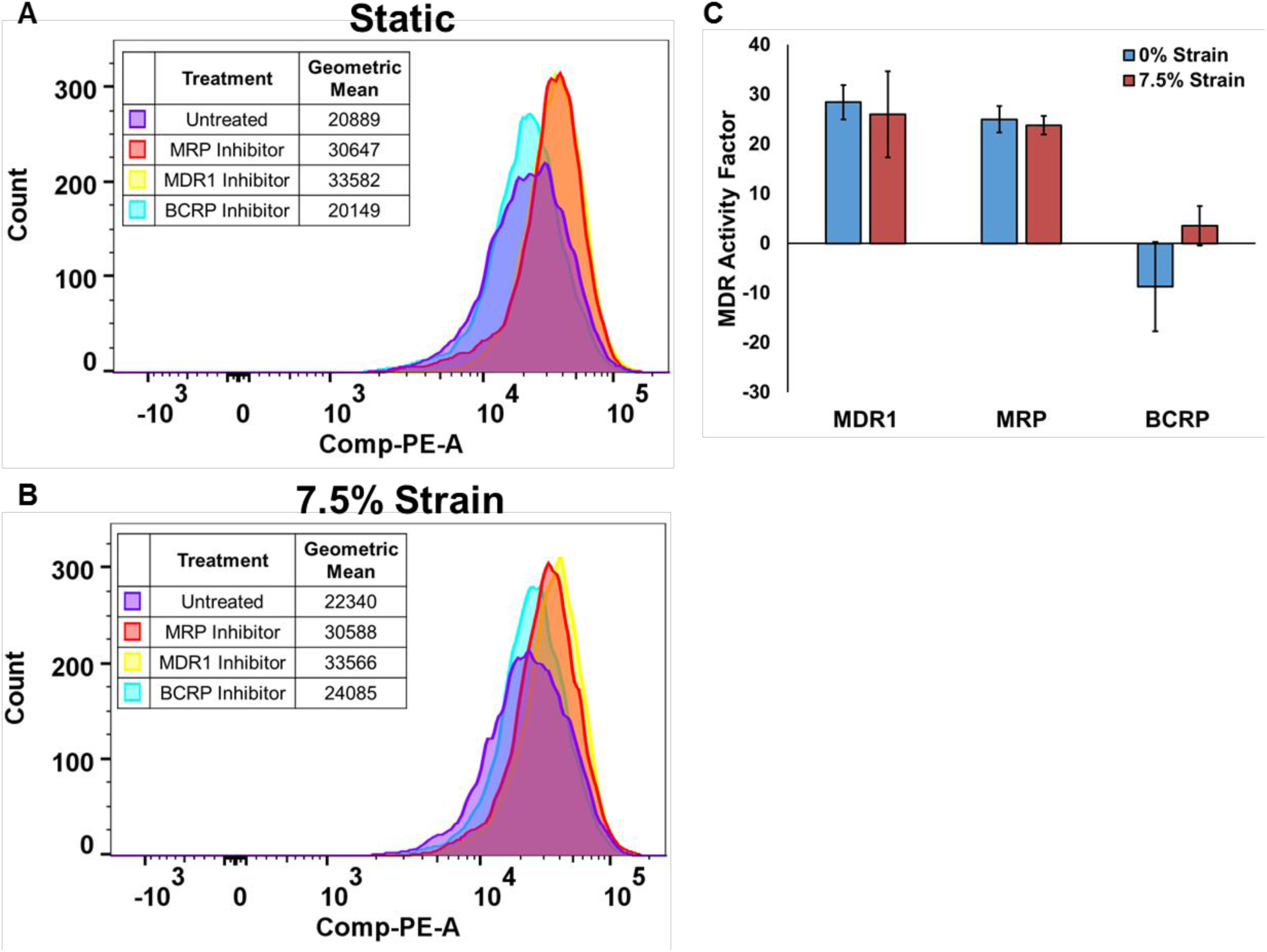
Mechanical strain does not induce changes in multidrug resistance protein activity in MDA-MB-231 breast cancer cells. MDA-MB-231 breast cancer cells were mechanically strained for 24 hours at 0 or 7.5% strain. The cells were then tested for activity of multidrug resistance proteins MDR1, MRP1, and BCRP. Flow cytometry histogram of cell counts for gold dye intensity in (A) statically cultured cells and (B) cells conditioned with 7.5% cyclic mechanical strain. Increasing presence of multidrug resistance efflux pumps would decrease concentration of gold dye inside the cell. (C) Multidrug resistance activity factor was calculated to determine the influence of three multidrug resistance efflux pumps, multi drug resistance protein 1 (MDR1), MDR-associated protein (MRP1), and breast cancer resistance protein (BCRP).

**Supplemental Figure 4.**
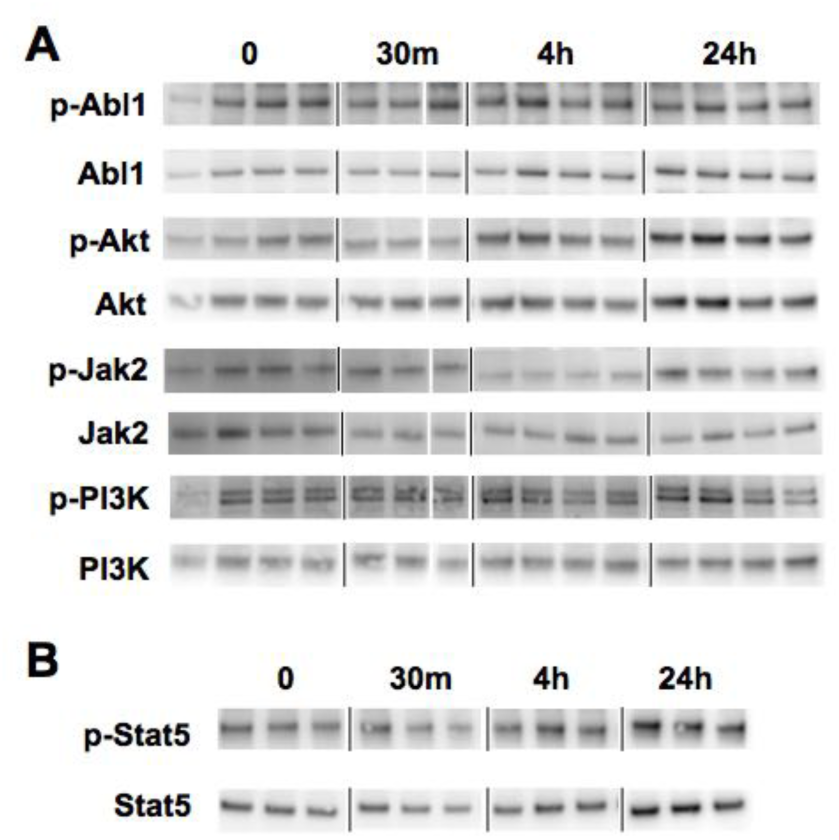
Summary diagram of mechanisms inferred by studies on mechanically induced enhancement in cell survival and chemoresistance.

**Supplemental Figure 5.**
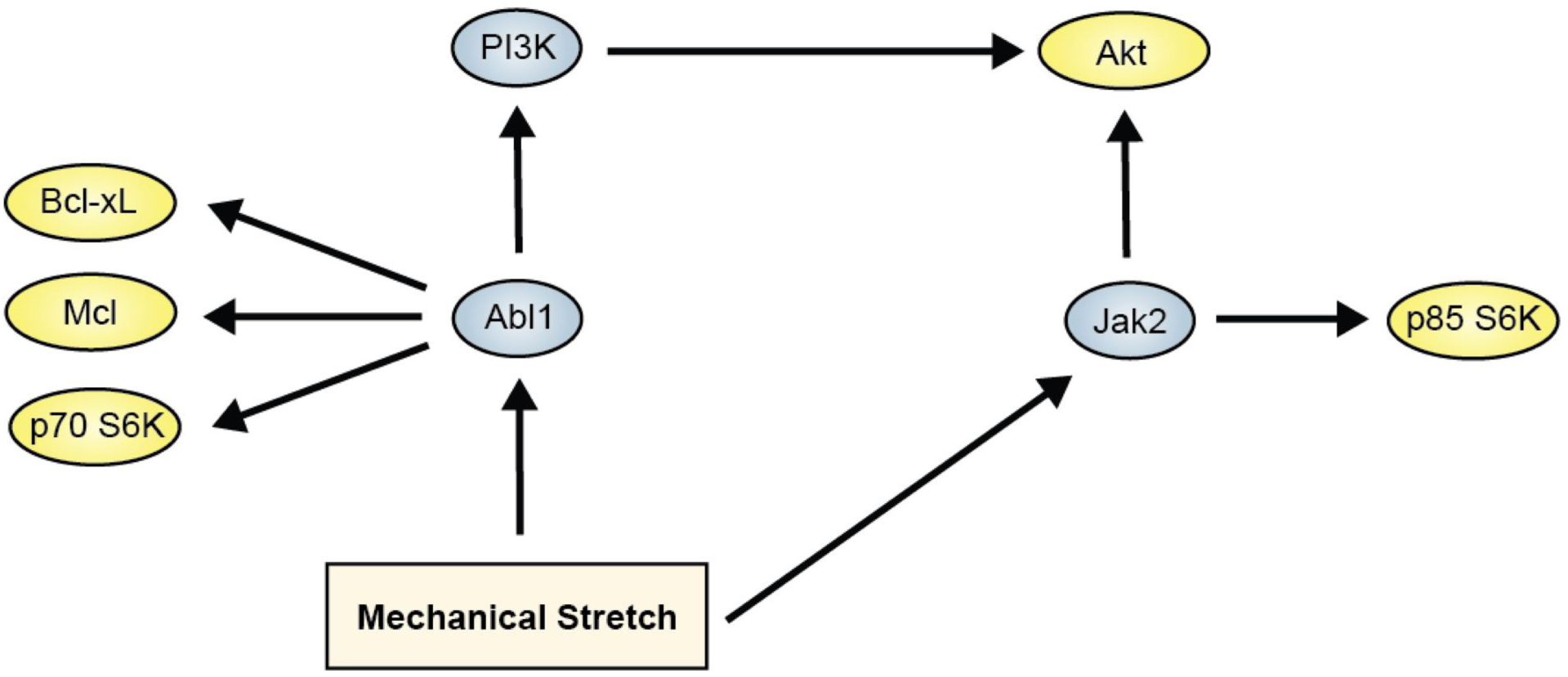
Vasculogenic activity is increased by mechanical loading in breast cancer cells. MBA-MB-231 cells were mechanically loaded for 24 hours at 7.5% strain at 1 Hz. (A) Endothelial cells were treated with conditioned media from mechanically loaded cancer cells in tube formation assay on matrigel. *p < 0.05 versus static group. (B) MDA-MB-231 cells were treated with mechanical load and then used directly in a tube formation assay on matrigel. *p < 0.05 versus static group.

**Supplemental Figure 6.**
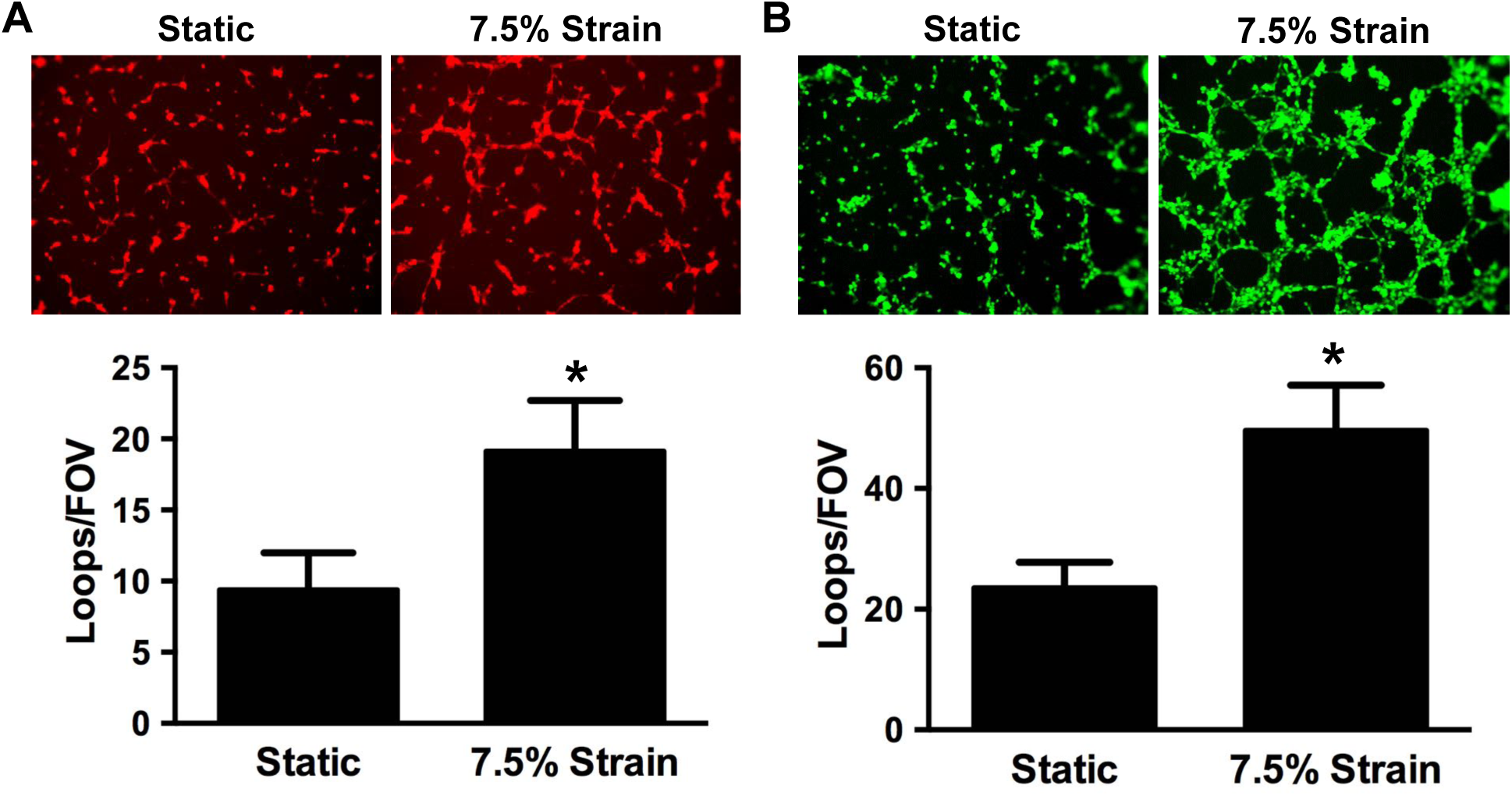
Tumor volumes for orthotopic implantation study. (A) Tumor volumes for tumors at three days. (B) Full time course of tumor volumes for the study.

**Supplemental Figure 7.**
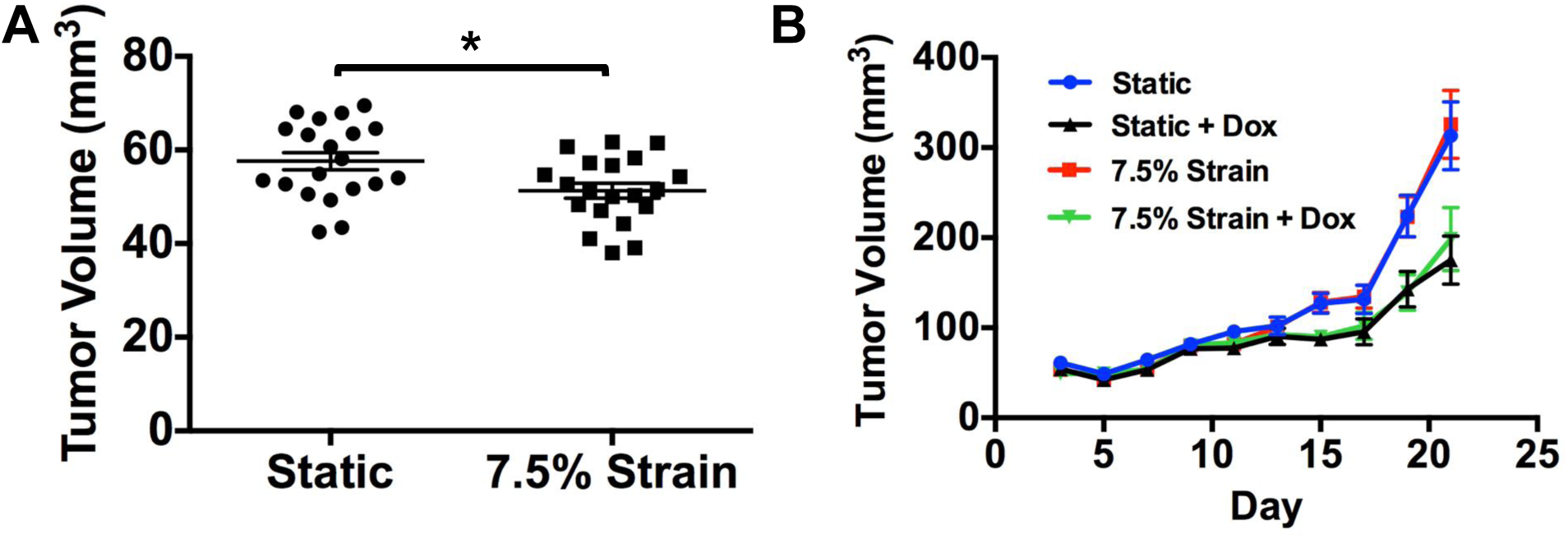
Tumor weights for excised tumors from the orthotopic implantation study.

**Supplemental Figure 8.**
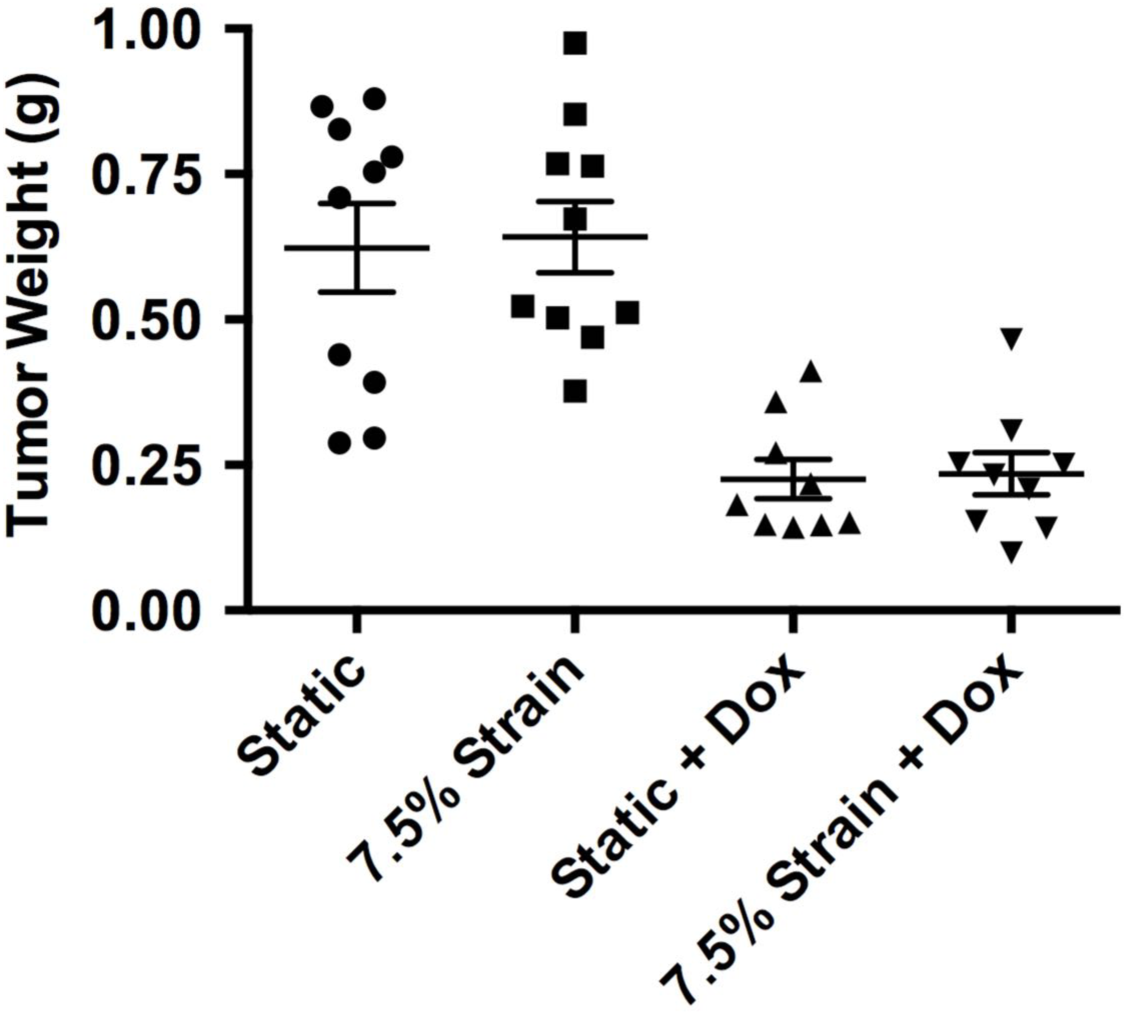
Laser speckle imaging for late time point in orthotopic tumor model. MDA-MB-231 cells were treated with mechanical load for 24 hours and then injected into the mammary fat pad of nu/nu mice. (A) Laser speckle images of the mice following treatment. The dashed circles illustrate the location of the mammary fat pad. The right mammary fat pad received the injection and the left served as a contralateral control. (B) Quantification of the relative perfusion of the injected fat pad to the non-injected fat pad.

**Supplemental Figure 9.**
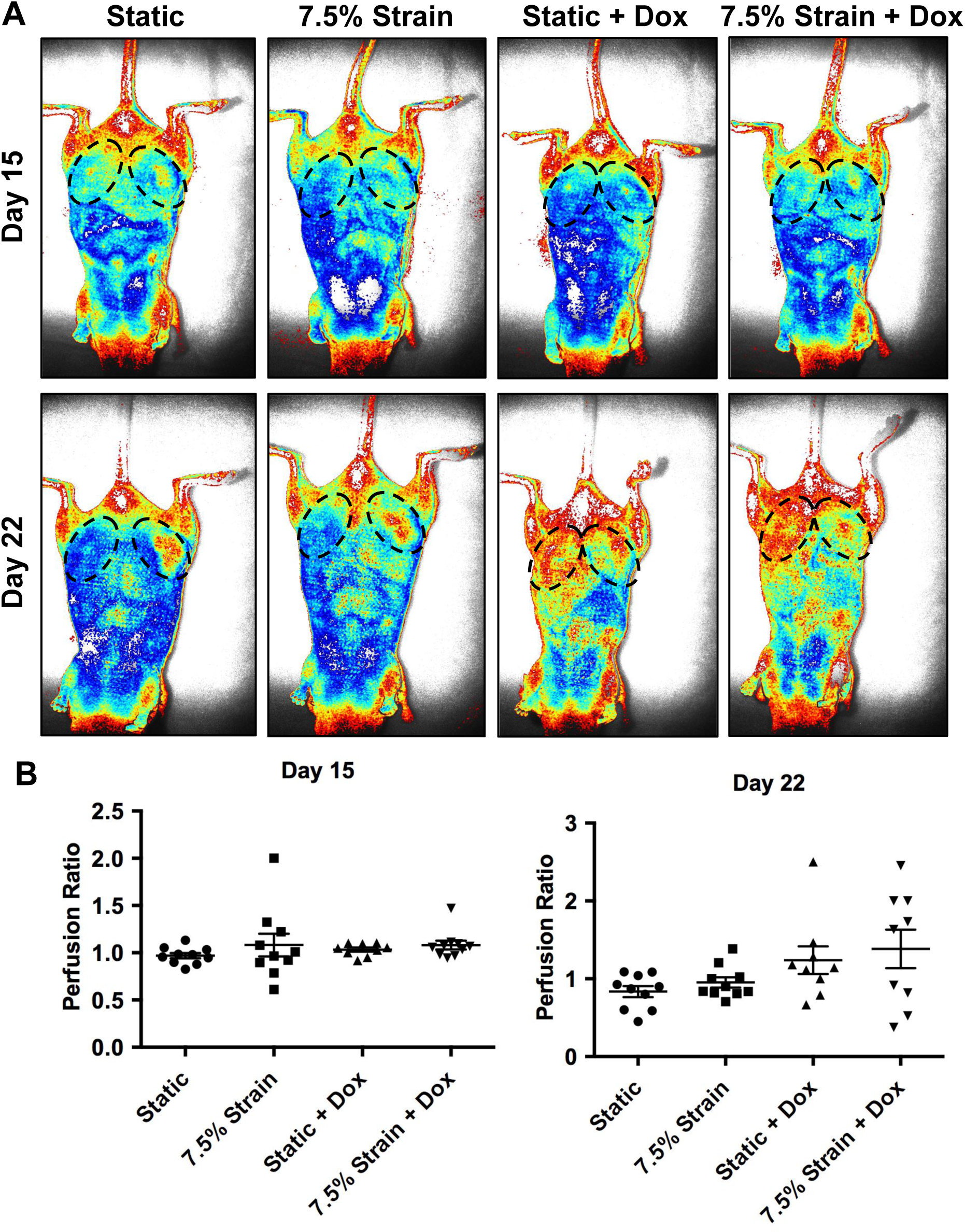
Micro-CT images of the skulls from mice injected with MDA-MB-231 cells grown under static conditions or 7.5% strain for 24 hours.

**Supplemental Figure 10.**
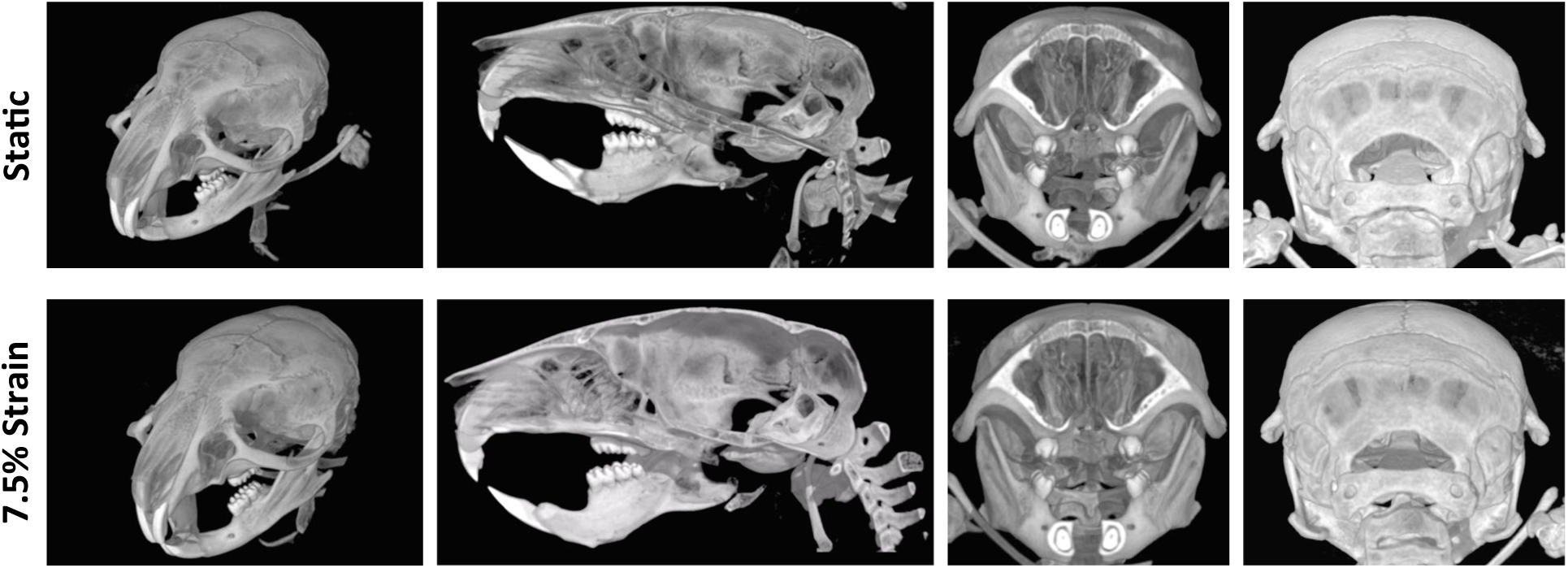

## Supplemental Tables

**Supplemental Table 1.**
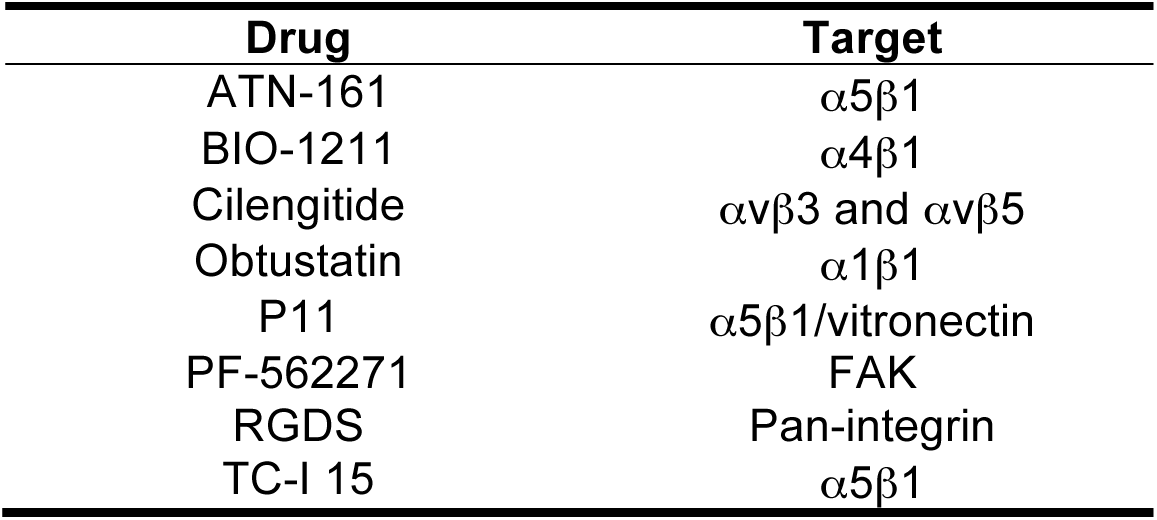
Integrin inhibitors Used in the Study.

**Supplemental Table 2.** Primary Antibodies/Reagents Used for Immunostaining

| **Target Protein** | **Company** | **Catalog #** | **Species/Isotype** | **Dilution Ratio** |
| --- | --- | --- | --- | --- |
| AF 594 Phalloidin | Invitrogen | A12381 | N/A | 1:250 |
| Paxillin | Santa Cruz Biotechnology | sc-5574 | anti-rabbit | 1:50 |
| PECAM-1 | Cell Signaling | 3528S | anti-mouse | 1:100 |
| α-SMA | Abcam | ab21027 | anti-rabbit | 1:100 |
| ILK | BD Biosciences | 611803 | anti-mouse | 1:100 |
| Yap/Taz | Cell Signaling | 8418S | anti-rabbit | 1:100 |
| 14-3-3ε | Santa Cruz Biotechnology | sc393177 | anti-mouse | 1:50 |
| Smad 2/3 | BD Biosciences | 610842 | anti-mouse | 1:100 |
| Phospho-Smad2/3 | Cell Signaling | 8828S | anti-rabbit | 1:100 |
| vWF | Santa Cruz Biotechnology | 365712 | anti-mouse | 1:50 |
| Myosin IIb | Cell Signaling | 3404 | anti-rabbit | 1:100 |
| Sca-1 | Santa Cruz Biotechnology | sc-365343 | anti-mouse | 1:50 |
| Oct-4 | Cell Signaling | 2750 | anti-rabbit | 1:100 |

**Supplemental Table 3.**
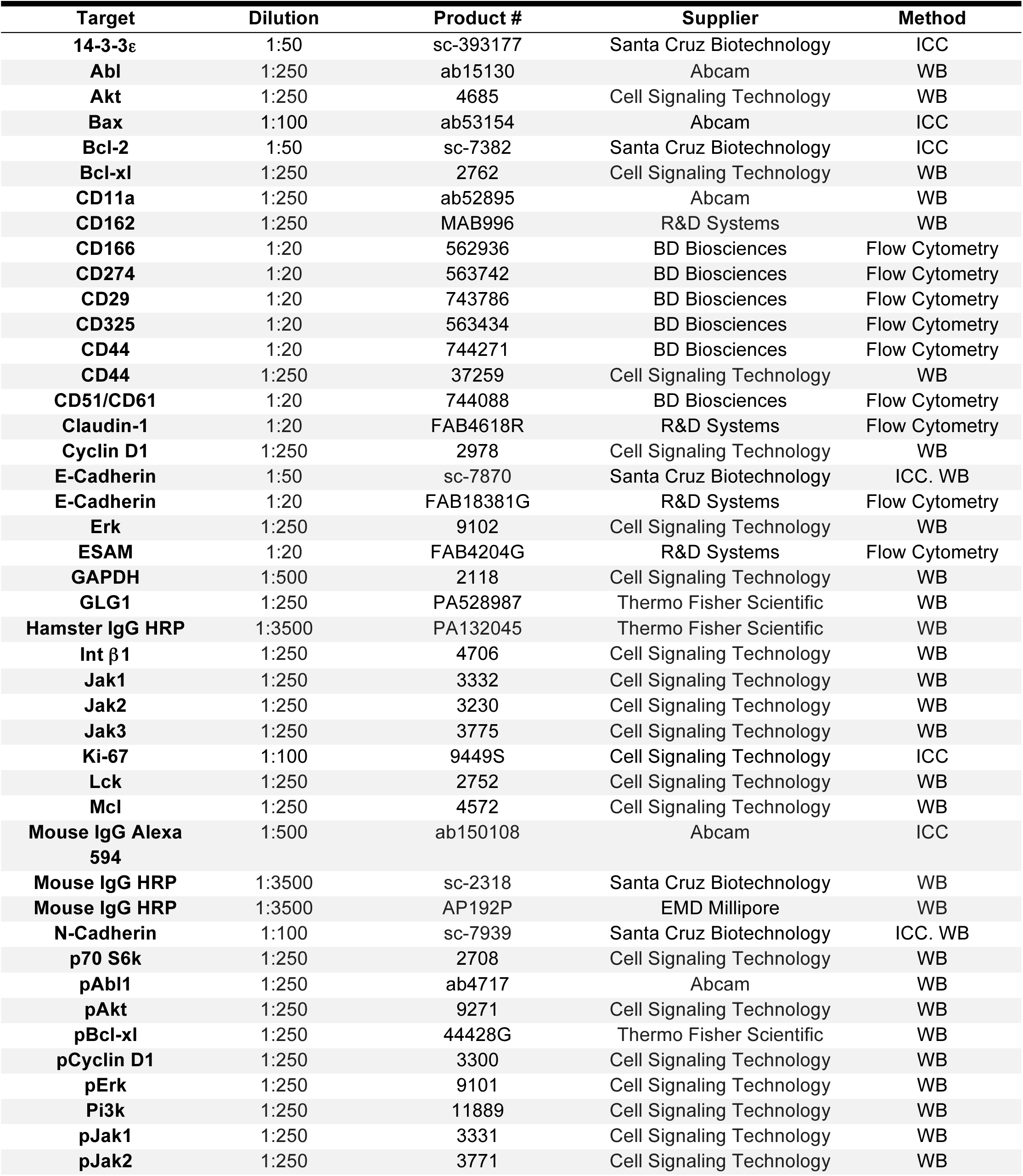
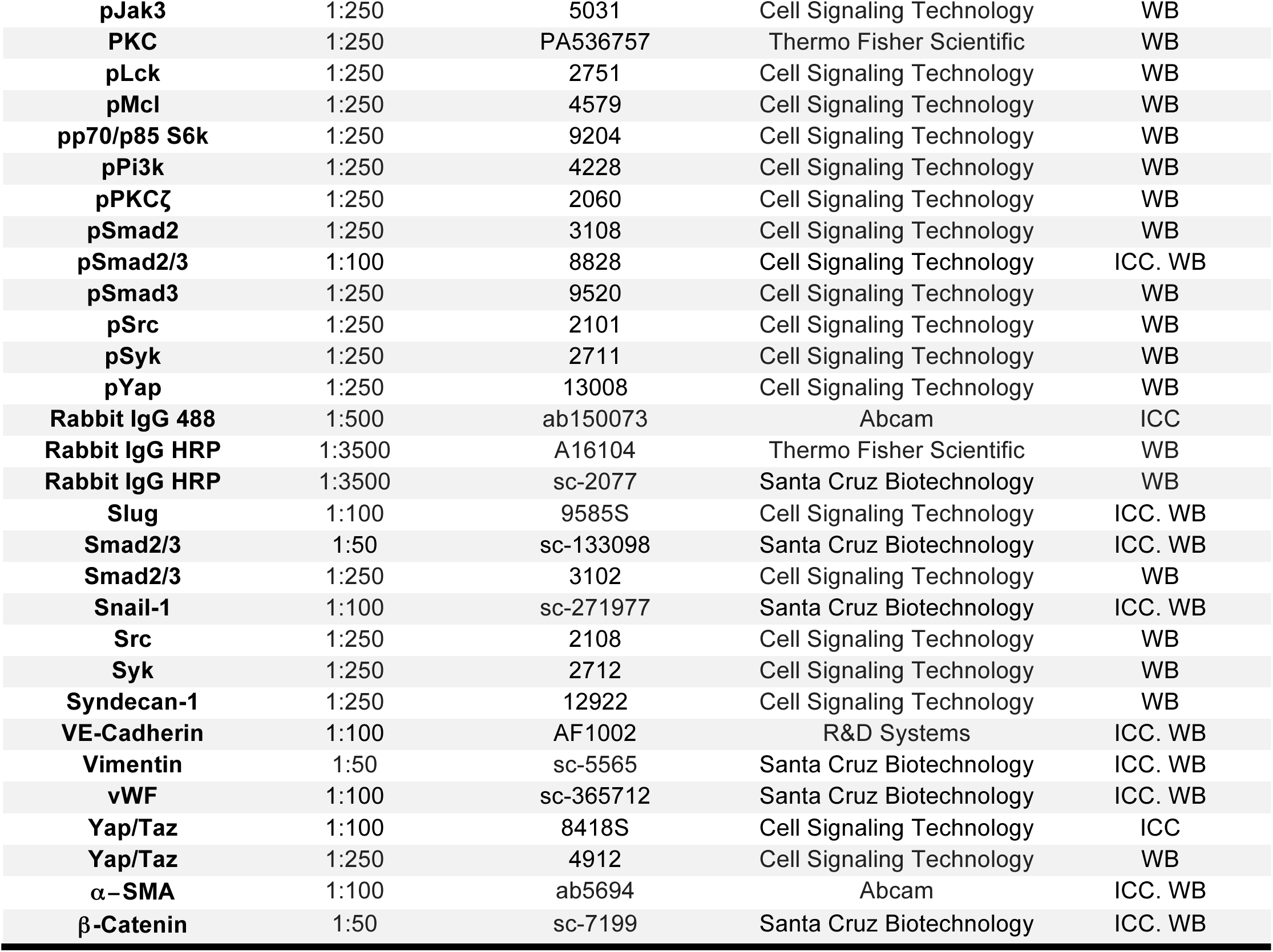
Antibodies Used in the Studies.

| **Target** | **Dilution** | **Product #** | **Supplier** | **Method** |
| --- | --- | --- | --- | --- |
| **14-3-3ε** | 1:50 | sc-393177 | Santa Cruz Biotechnology | ICC |
| **Abl** | 1:250 | ab15130 | Abcam | WB |
| **Akt** | 1:250 | 4685 | Cell Signaling Technology | WB |
| **Bax** | 1:100 | ab53154 | Abcam | ICC |
| **Bcl-2** | 1:50 | sc-7382 | Santa Cruz Biotechnology | ICC |
| **Bcl-xl** | 1:250 | 2762 | Cell Signaling Technology | WB |
| **CD11a** | 1:250 | ab52895 | Abcam | WB |
| **CD162** | 1:250 | MAB996 | R&D Systems | WB |
| **CD166** | 1:20 | 562936 | BD Biosciences | Flow Cytometry |
| **CD274** | 1:20 | 563742 | BD Biosciences | Flow Cytometry |
| **CD29** | 1:20 | 743786 | BD Biosciences | Flow Cytometry |
| **CD325** | 1:20 | 563434 | BD Biosciences | Flow Cytometry |
| **CD44** | 1:20 | 744271 | BD Biosciences | Flow Cytometry |
| **CD44** | 1:250 | 37259 | Cell Signaling Technology | WB |
| **CD51/CD61** | 1:20 | 744088 | BD Biosciences | Flow Cytometry |
| **Claudin-1** | 1:20 | FAB4618R | R&D Systems | Flow Cytometry |
| **Cyclin D1** | 1:250 | 2978 | Cell Signaling Technology | WB |
| **E-Cadherin** | 1:50 | sc-7870 | Santa Cruz Biotechnology | ICC. WB |
| **E-Cadherin** | 1:20 | FAB18381G | R&D Systems | Flow Cytometry |
| **Erk** | 1:250 | 9102 | Cell Signaling Technology | WB |
| **ESAM** | 1:20 | FAB4204G | R&D Systems | Flow Cytometry |
| **GAPDH** | 1:500 | 2118 | Cell Signaling Technology | WB |
| **GLG1** | 1:250 | PA528987 | Thermo Fisher Scientific | WB |
| **Hamster IgG HRP** | 1:3500 | PA132045 | Thermo Fisher Scientific | WB |
| **Int β1** | 1:250 | 4706 | Cell Signaling Technology | WB |
| **Jak1** | 1:250 | 3332 | Cell Signaling Technology | WB |
| **Jak2** | 1:250 | 3230 | Cell Signaling Technology | WB |
| **Jak3** | 1:250 | 3775 | Cell Signaling Technology | WB |
| **Ki-67** | 1:100 | 9449S | Cell Signaling Technology | ICC |
| **Lck** | 1:250 | 2752 | Cell Signaling Technology | WB |
| **Mcl** | 1:250 | 4572 | Cell Signaling Technology | WB |
| **Mouse IgG Alexa 594** | 1:500 | ab150108 | Abcam | ICC |
| **Mouse IgG HRP** | 1:3500 | sc-2318 | Santa Cruz Biotechnology | WB |
| **Mouse IgG HRP** | 1:3500 | AP192P | EMD Millipore | WB |
| **N-Cadherin** | 1:100 | sc-7939 | Santa Cruz Biotechnology | ICC. WB |
| **p70 S6k** | 1:250 | 2708 | Cell Signaling Technology | WB |
| **pAbl1** | 1:250 | ab4717 | Abcam | WB |
| **pAkt** | 1:250 | 9271 | Cell Signaling Technology | WB |
| **pBcl-xl** | 1:250 | 44428G | Thermo Fisher Scientific | WB |
| **pCyclin D1** | 1:250 | 3300 | Cell Signaling Technology | WB |
| **pErk** | 1:250 | 9101 | Cell Signaling Technology | WB |
| **Pi3k** | 1:250 | 11889 | Cell Signaling Technology | WB |
| **pJak1** | 1:250 | 3331 | Cell Signaling Technology | WB |
| **pJak2** | 1:250 | 3771 | Cell Signaling Technology | WB |
| **pJak3** | 1:250 | 5031 | Cell Signaling Technology | WB |
| **PKC** | 1:250 | PA536757 | Thermo Fisher Scientific | WB |
| **pLck** | 1:250 | 2751 | Cell Signaling Technology | WB |
| **pMcl** | 1:250 | 4579 | Cell Signaling Technology | WB |
| **pp70/p85 S6k** | 1:250 | 9204 | Cell Signaling Technology | WB |
| **pPi3k** | 1:250 | 4228 | Cell Signaling Technology | WB |
| **pPKCζ** | 1:250 | 2060 | Cell Signaling Technology | WB |
| **pSmad2** | 1:250 | 3108 | Cell Signaling Technology | WB |
| **pSmad2/3** | 1:100 | 8828 | Cell Signaling Technology | ICC. WB |
| **pSmad3** | 1:250 | 9520 | Cell Signaling Technology | WB |
| **pSrc** | 1:250 | 2101 | Cell Signaling Technology | WB |
| **pSyk** | 1:250 | 2711 | Cell Signaling Technology | WB |
| **pYap** | 1:250 | 13008 | Cell Signaling Technology | WB |
| **Rabbit IgG 488** | 1:500 | ab150073 | Abcam | ICC |
| **Rabbit IgG HRP** | 1:3500 | A16104 | Thermo Fisher Scientific | WB |
| **Rabbit IgG HRP** | 1:3500 | sc-2077 | Santa Cruz Biotechnology | WB |
| **Slug** | 1:100 | 9585S | Cell Signaling Technology | ICC. WB |
| **Smad2/3** | 1:50 | sc-133098 | Santa Cruz Biotechnology | ICC. WB |
| **Smad2/3** | 1:250 | 3102 | Cell Signaling Technology | WB |
| **Snail-1** | 1:100 | sc-271977 | Santa Cruz Biotechnology | ICC. WB |
| **Src** | 1:250 | 2108 | Cell Signaling Technology | WB |
| **Syk** | 1:250 | 2712 | Cell Signaling Technology | WB |
| **Syndecan-1** | 1:250 | 12922 | Cell Signaling Technology | WB |
| **VE-Cadherin** | 1:100 | AF1002 | R&D Systems | ICC. WB |
| **Vimentin** | 1:50 | sc-5565 | Santa Cruz Biotechnology | ICC. WB |
| **vWF** | 1:100 | sc-365712 | Santa Cruz Biotechnology | ICC. WB |
| **Yap/Taz** | 1:100 | 8418S | Cell Signaling Technology | ICC |
| **Yap/Taz** | 1:250 | 4912 | Cell Signaling Technology | WB |
| **α-SMA** | 1:100 | ab5694 | Abcam | ICC. WB |
| **β-Catenin** | 1:50 | sc-7199 | Santa Cruz Biotechnology | ICC. WB |

